# Cell-free, methylated DNA in blood samples reveals tissue-specific, cellular damage from radiation treatment

**DOI:** 10.1101/2022.04.12.487966

**Authors:** Megan E. Barefoot, Netanel Loyfer, Amber J. Kiliti, Marcel O. Schmidt, Sapir Shabi-Porat, Sidharth Jain, Sarah Martinez Roth, A. Patrick McDeed, Nesreen Shahrour, Elizabeth Ballew, Yun-Tien Lin, Heng-Hong Li, Anne Deslattes Mays, Sonali Rudra, Anna T. Riegel, Keith Unger, Tommy Kaplan, Anton Wellstein

## Abstract

Radiation therapy is an effective cancer treatment although damage to healthy tissues is common. Here we characterize the methylomes of healthy human and mouse tissues to establish sequencing-based, cell-type specific reference DNA methylation atlases. Identified cell-type specific DNA blocks were mostly hypomethylated and located within genes intrinsic to cellular identity. Cell-free DNA fragments released from dying cells into the circulation were captured from serum samples by hybridization to CpG-rich DNA panels. The origins of the circulating DNA fragments were inferred from mapping to the established DNA methylation atlases. Thoracic radiation-induced tissue damages in a mouse model were reflected by dose-dependent increases in lung endothelial, cardiomyocyte and hepatocyte methylated DNA in serum. The analysis of serum samples from breast cancer patients undergoing radiation treatment revealed distinct tissue-specific epithelial and endothelial responses to radiation across multiple organs. Strikingly, patients treated for right-sided breast cancers also showed increased hepatocyte and liver endothelial DNA in the circulation indicating the impact on liver tissues. Thus, changes in cell-free methylated DNA can uncover cell-type specific effects of radiation and provide a quantitative measure of the biologically effective radiation dose received by healthy tissues.

**Graphical Abstract:** 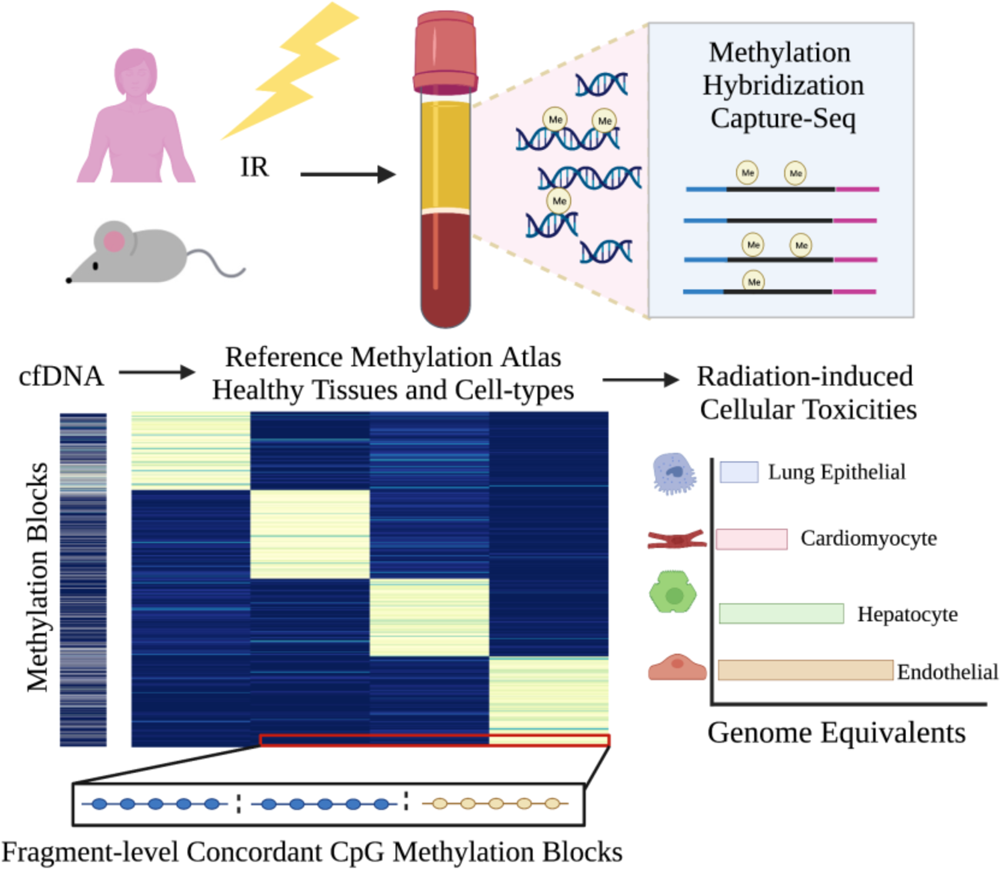

## Introduction

Radiation therapy is an effective cancer treatment; however, surrounding normal tissues are also impacted leading to tissue damage and remodeling(1–4). For breast cancer patients, the heart, lungs, and skin are the most common organs-at-risk for toxicity(5–7). However, radiation-induced toxicities vary due to patient-specific factors and clinical symptoms may be acute or long-term, often appearing months or even years after treatment(6). The mechanisms of radiation-induced tissue injury are still poorly characterized and there are few biomarkers of radiation-related damage. Here we address this unmet need for sensitive and tissue-specific detection of cellular injury using serially collected blood samples.

Decoding the cellular origins of circulating cell-free DNA (cfDNA) from blood samples (“liquid biopsies”) is a promising approach for non-invasive monitoring of organ homeostasis, where rising levels of cfDNA released from dying cells indicate increased tissue damage(8–13). The majority of cfDNA fragments peak around 167 bp, corresponding to the length of DNA wrapped around a nucleosome (147 bp) plus a linker fragment (20 bp). This nucleosomal footprint in cfDNA reflects degradation by nucleases as a by-product of cell death and the tissue origins of the cfDNA fragments can be uncovered using highly cell-type specific DNA methylation patterns(8, 14). DNA methylation typically involves covalent addition of a methyl group to the 5-carbon of cytosine (5mc) with the human and mouse genomes containing 28 and 13 million CpG sites respectively(14, 15). Dynamic changes to the methylome during development and cellular differentiation lead to stable, cell-type specific patterns of DNA methylation that are conserved during DNA replication and thus provide the predominant mechanism for inherited cellular memory during cell growth(16–19).

Recent studies have demonstrated the feasibility of Tissue-Of-Origin (TOO) analysis in the circulation using cell-free DNA methylation(20–24). However, few of these studies have focused on tracking intervention-related changes over time that is feasible by analyses of serially collected liquid biopsies(25–27). The short half-life of cfDNA (15 mins – 2 hours) is ideal for detecting real-time changes in tissue homeostasis due to therapeutic interventions(27, 28). Also, few cfDNA analyses have taken advantage of CpG pattern analysis to increase sensitivity and specificity of cell type proportion estimates(22, 23, 29–32). Each cfDNA molecule originates from a defined cell and pattern analysis of sequence reads allows for individual classification of each sequenced fragment as opposed to traditional methods that average the methylation status across a population of fragments aligned at single CpG sites(27, 28). Building on these advances, we present a fine-tuned approach for deconvolution of cfDNA patterns based on fragment-level CpG methylation blocks.

Here, we first report comprehensive, sequencing-based DNA methylation reference maps of healthy human and mouse cell-types and show the close relationship of DNA methylation with cellular gene expression. Then, we apply cell-type specific DNA methylation to trace the origins of cfDNA in serum samples. We report that hybridization capture sequencing of methylated cfDNA in serum samples reflects dose-dependent tissue damages in a mouse model of radiation injury. In addition, analyses of serial serum samples from breast cancer patients undergoing routine radiation treatment indicate distinct cellular damages in different organs providing a measure of the biologically effective radiation doses being administered. As proof of concept, radiation treatment serves as a powerful tool to validate the cell-type specific methylation signatures developed in the atlases and demonstrates application to detect cellular injury in the circulation.

## Results

### Experimental paradigm to identify the cellular origins of radiation-induced damage from cfDNA in the circulation

To investigate if radiation-induced tissue damages can be monitored from changes in methylated cfDNA in the circulation, we collected serial serum samples from breast cancer patients undergoing routine radiation treatment as well as serum and tissue samples from mice that had received different doses of thoracic radiation (**Figure 1**). The bioanalyzer trace in **Figure 1** shows readings of cfDNA isolated from serum samples that were bisulfite treated, enriched for sequences of interest by methylome-wide hybridization capture and subjected to sequence analysis. As a prerequisite for identification of the cellular origins of the cfDNA fragments isolated from the circulation, we established human and mouse cell-type specific DNA methylation atlases. We took a sequencing-based approach interrogating existing WGBS data sets and generated complementary data from additional cell-types composing at-risk organs that include the lungs, heart, and liver (**Supplemental Table S1 and S2**). The characteristics and validation of the DNA methylation blocks that provide the basis for the cell-type specific mouse and human atlases are described next.

**Figure 1.**
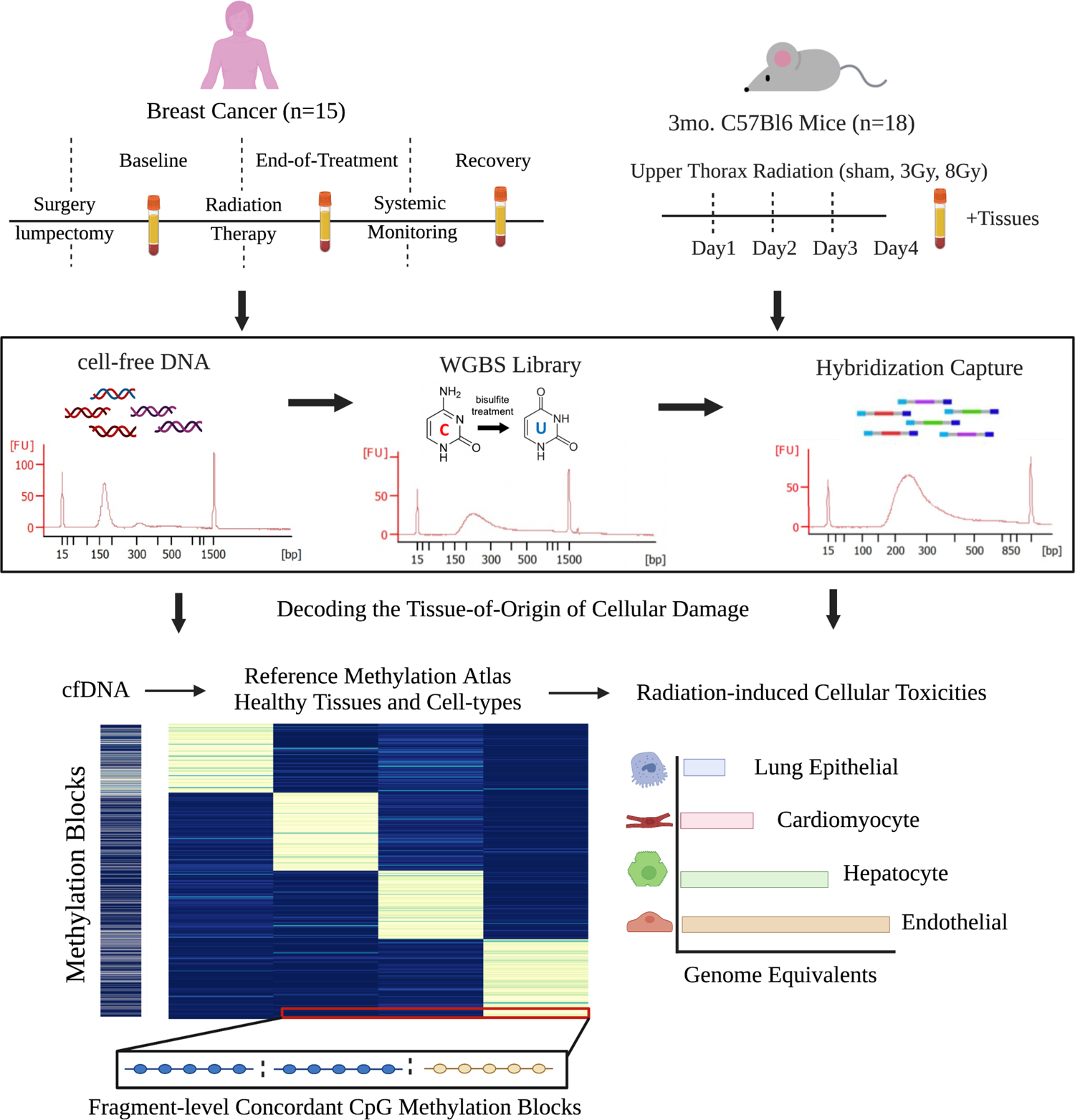
Experimental paradigm using cell-free methylated DNA in blood to identify cellular origins of radiation-induced tissue damages. Serial serum samples were collected from breast cancer patients treated with radiation. In parallel, paired serum and tissue samples were collected from mice receiving radiation at 3Gy or 8Gy doses compared to sham control. Cell-free DNA (cfDNA) methylome profiling of serum samples was performed using hybridization capture-sequencing of bisulfite-treated cfDNA. Cell-type specific methylation blocks were identified from Whole Genome Bisulfite Sequencing (WGBS) reference data of healthy tissues and used to identify the cellular origins of the serum cfDNA.

### Differences in DNA methylation blocks reflect distinct developmental lineages and cellular identities

We obtained controlled access to reference human and mouse WGBS datasets from publicly available databases, preferentially from primary cells isolated from healthy human and mouse tissues. Additionally, we generated cell-type specific methylomes for mouse immune cell types (CD19+ B cells, Gr1+ Neutrophils, CD4+ T cells, and CD8+ T cells) and human tissue-specific endothelial cell types (coronary artery, pulmonary artery, cardiac microvascular and liver sinusoidal endothelial cells). This resulted in curation of mouse methylation data from 10 different cell types and 18 tissues to establish the most comprehensive mouse methylation atlas to date. In addition, we characterized methylation data from over 30 distinct human cell types with a diverse representation of donors (**Supplemental Tables 1 and 2**). To better understand the epigenomic landscape of these healthy human and mouse cell types in tissues, we characterized the methylomes by first segmenting the data into homogenously methylated blocks where DNA methylation levels at adjacent CpG sites are highly correlated across different cell types(22). Curated human WGBS datasets from healthy cell types were segmented to identify 351,395 blocks covered by our hybridization capture panel used in the analysis of cfDNA in human serum (captures 80Mb, ∼20% of CpGs). Likewise, segmentation of mouse WGBS datasets from healthy cell types and tissues identified 1,344,889 blocks covered by the mouse hybridization capture panel (captures 210 Mb, ∼75% of CpGs). Unsupervised hierarchical clustering analysis of the top 30,000 variable methylation blocks amongst all human samples revealed that cellular identity and developmental lineage primarily drives the relationship between samples and is presented as a dendrogram and UMAP projection in **Figure 2a and 2b**. The respective analysis of mouse cell types is depicted in **Figure 2c** and **Supplemental Figure 1**. The tight relationship of methylomes of the same cell type observed from the cluster analysis reinforces the concept that methylation status is conserved at regions critical to cell identity. The variation in distance between all samples was ∼12x larger than the variation in distance between samples from the same cell type. This stability allows methylated DNA to serve as a robust biomarker of cell types across diverse populations. For the most part, cells composing distinct lineages remain closely related, including immune, epithelial, muscle, neuron, endothelial, and stromal cell types. Examples are tissue-specific endothelial and tissue-resident immune cells that cluster with endothelial or immune cells respectively, independent of the germ layer origin of their tissues of residence. Collectively, these findings support that DNA methylation is highly cell-type specific and reflects cell lineage specification.

**Figure 2.**
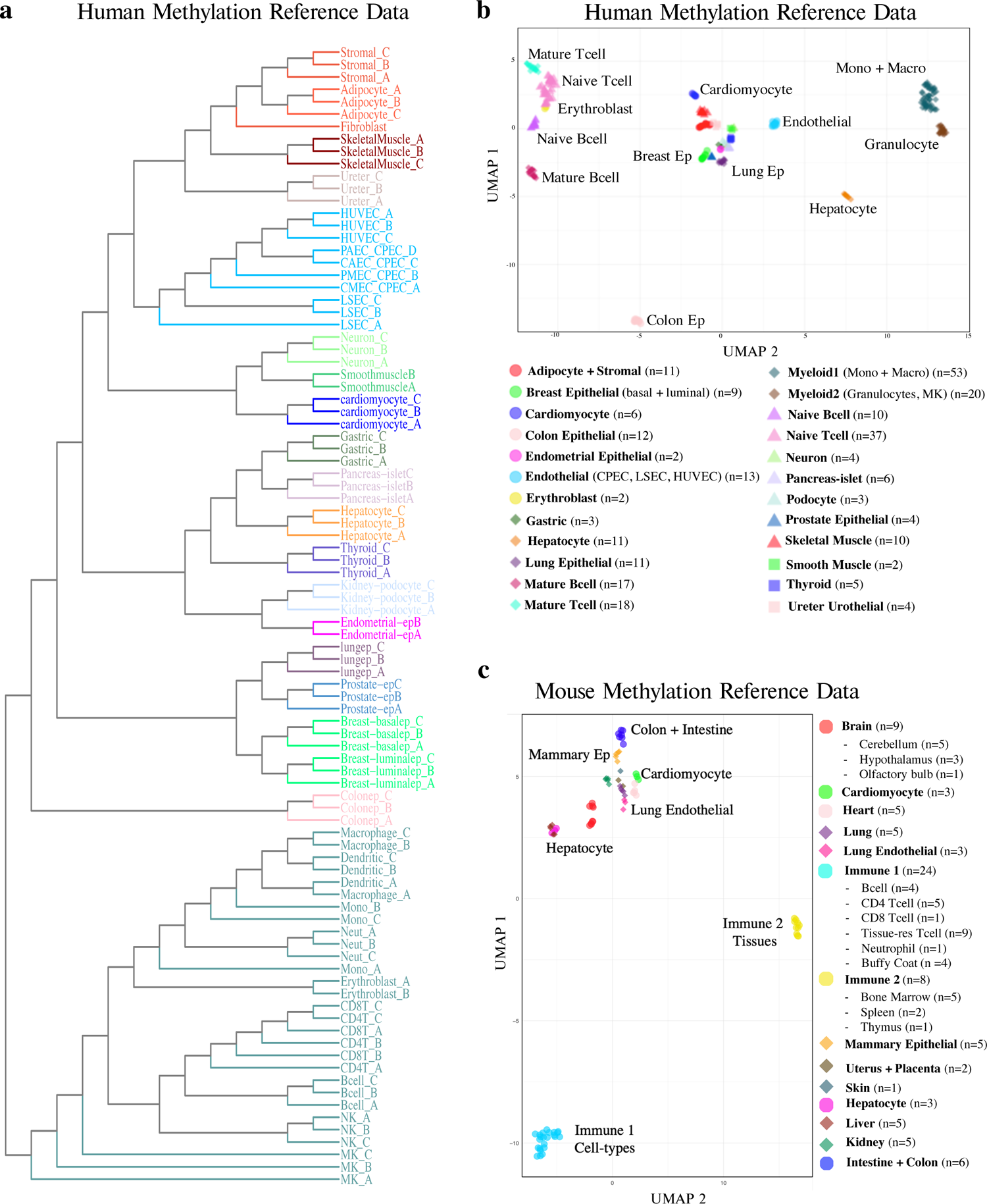
Characterization of human and mouse cell-type specific reference methylation data. (a) Tree dendrogram depicting the relationship between different cell types. Whole Genome Bisulfite Sequencing (WGBS) datasets were included in the analysis. Methylation status of the top 30,000 variable blocks was used as input for the unsupervised hierarchical clustering. Samples from cell types with greater than n=3 replicates were merged. (b, c) UMAP projection of human (b) and mouse (c) WGBS reference datasets. Abbreviations: CAEC = coronary artery endothelial cell, CMEC = cardiac microvascular endothelial cell, CPEC = joint cardio-pulmonary endothelial cell, HUVEV = human umbilical vein endothelial cell, LSEC = liver sinusoidal endothelial cell, MK = megakaryocyte, NK = natural killer cell, PAEC = pulmonary artery endothelial cell, PMEC = pulmonary microvascular endothelial cell.

### Development of sequencing-based DNA methylation atlases of primary human and mouse cell-types

Based on the above unsupervised clustering analysis, we selected a final set of reference methylomes used to identify differentially methylated cell-type specific blocks. We excluded WGBS samples from bulk tissues and samples with low coverage. Subsets of some related cell types were considered together to form the final groups (i.e., monocytes grouped together with macrophages and colon grouped together with small intestine). We identified cell-type specific differentially methylated blocks (DMBs) that contained a minimum of 3 CpG sites and overlapped with captured regions from our hybridization panels used in the analysis of cfDNA from serum. The co-methylation status of neighboring CpG sites in these blocks distinguished amongst all cell types included in the final groups. Overall, we identified 4,637 human and 7,344 mouse DMBs with a methylation difference of 0.35 for mouse and 0.4 for human. The human and mouse DNA methylation blocks specific for these cell types can be found in **Supplemental Tables 3 and 4.** A summary of human and mouse cell-type specific methylation blocks is in **Supplemental Table 5**. Intriguingly, a variable number of blocks were identified for each cell-type using the same thresholds. This is likely due to genuine biologic differences between cell types but also impacted by the depth of coverage, purity, and degree of separation from other tissues and cell types currently included in the atlas. To visualize cell-type specific DMBs, we created a methylation score that applies to both hypomethylated and hypermethylated DMBs. The score calculates the number of fully unmethylated read-pairs divided by total coverage for hypomethylated blocks (and vice versa for hypermethylated blocks). The heatmaps in **Figure 3a and 3b** depict up to 100 blocks with the highest methylation score for each cell-type group.

**Figure 3.**
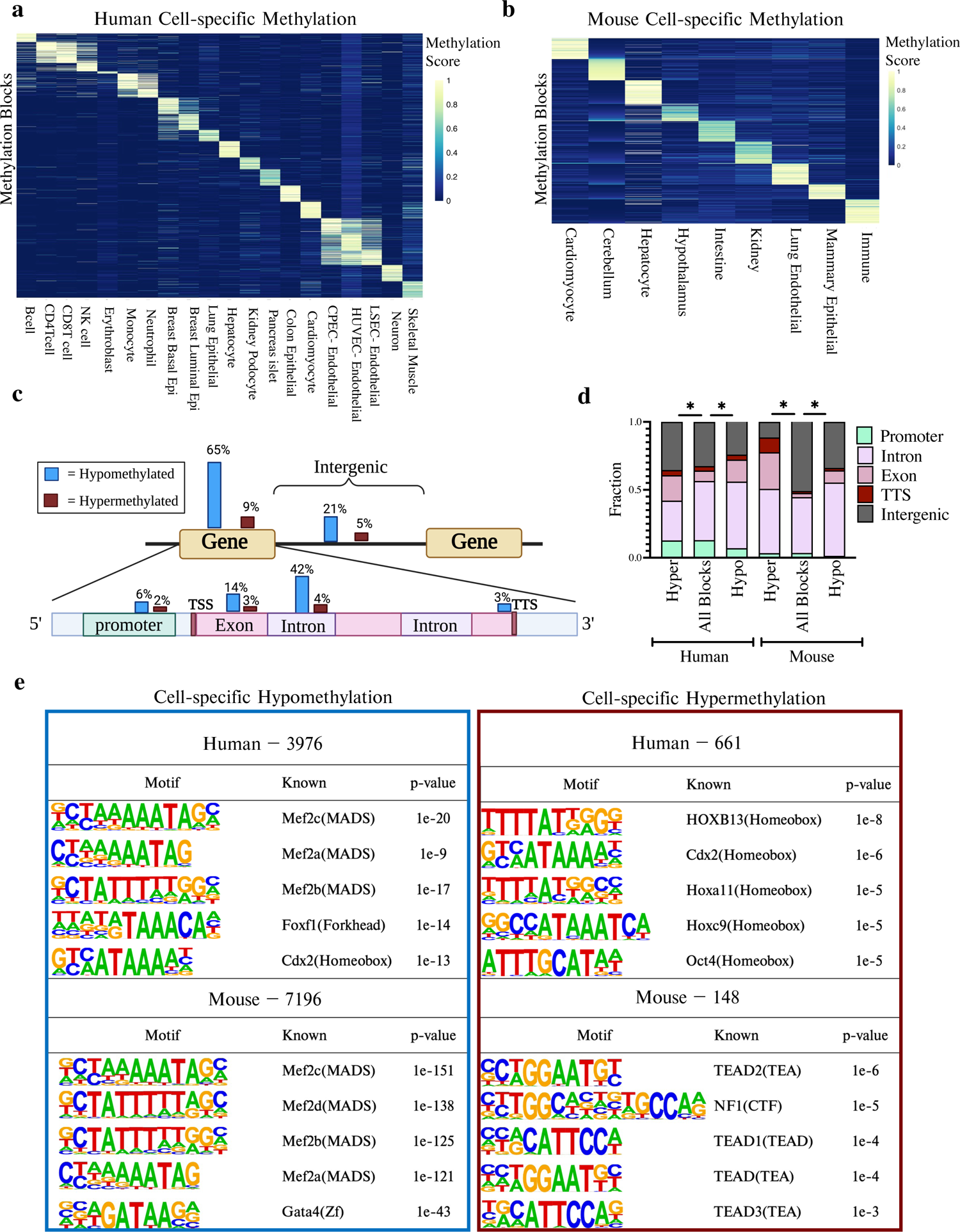
Cell-type specific DNA blocks are mostly hypomethylated, enriched at intragenic regions and developmental transcription factor (TF) binding motifs. (a-b) Heatmaps of differentially methylated cell-type specific blocks identified from reference WGBS data compiled from healthy cell types and tissues in human (a) and mouse (b). Each cell in the plot marks the methylation score of one genomic region (rows) at each of the 20 cell types in human and 9 in mouse (columns). Up to 100 blocks with the highest methylation score are shown per cell type. The methylation score represents the number of fully unmethylated or methylated read-pairs / total coverage for hypo- and hyper-methylated blocks, respectively. (c) Schematic diagram depicting location of human cell-type specific hypo- and hyper-methylated blocks. Genomic annotations of cell-type specific methylation blocks were determined by analysis using HOMER. (d) Distribution of human (left) and mouse (right) cell-type specific methylation blocks relative to genomic regions used in the hybridization capture probes. Captured blocks with less than 5% variance across cell types represent blocks without cell-type specificity and were used as background. (e) Top 5 TF binding sites enriched among cell-type specific hypo- and hyper-methylated blocks in human (top) and mouse (bottom), using HOMER motif analysis. As above, captured blocks with less than 5% variance across cell types were used as background.

### Differential DNA methylation is closely linked to the regulation of cell-type specific functions

We next sought to understand the role of cell-type specific methylation in shaping cellular identity and function. For this we identified genes adjacent to cell-type specific methylation blocks and performed pathway analysis of annotated genes using both Ingenuity Pathway Analysis (IPA) and GREAT(42, 43). Important biological differences were also observed in the gene sets identified based on specific processes unique to the cell types profiled.

For example, the biological function of genes associated with immune cell-type specific methylation reflects processes of leukocyte cell-cell adhesion, immune response-regulating signaling, and hematopoietic system development (**Supplemental Figure 2**). In contrast, fatty acid metabolic process, lipid metabolism, and acute phase response signaling were identified for hepatocytes. Significantly enriched biological pathways and functions for genes associated with differential methylation in each cell type examined are provided in **Supplemental Table 11**.

### Cell-type specific DNA blocks are mostly hypomethylated and enriched at intragenic regions containing developmental TF binding motifs

The majority of human and mouse cell-type specific blocks identified here were hypomethylated, consistent with other studies(14, 17). In human samples we found 86% of cell-type specific DMBs hypomethylated and only 14% hypermethylated. Strikingly, in the mouse samples, 98% of cell-type specific DMBs were hypomethylated and only 2% were hypermethylated. The schematic in **Figure 3c** depicts the location of identified human cell-type specific hypo- and hyper-methylated blocks. Interestingly, regardless of directionality the majority of cell-type specific blocks were located within intragenic regions. To assess if the genomic locations of cell-type specific blocks are distinct, we compared the locations to the captured blocks that do not vary amongst cell types (**Figure 3d; Supplemental Table 10**). We found that for both human and mouse, there was a significant enrichment of cell-type specific blocks within intragenic regions relative to other captured regions (Fisher’s exact test, p<0.05). There was also a significant relationship between directionality and intragenic distribution, with a significantly larger proportion of cell-type specific blocks being hypermethylated in exons and hypomethylated in introns (Chi-square, df=3, p<0.05). The similar distribution of cell-type specific methylation blocks in human and mouse suggests a conserved biological function of these genomic regions across species.

To further explore what common function these identified regions may have in human and mouse development, we performed motif analysis using HOMER to see if there were commonly enriched transcription factor binding sites (TFBS)(41). MADS motifs bound by MEF2 transcription factors were significantly enriched in both human and mouse cell-type specific hypomethylated blocks (**Figure 3e - left)**. The MEF2 transcription factors are established developmental regulators with roles in the differentiation of many cell types from distinct lineages. In contrast, Homeobox motifs bound by several different HOX TFs were enriched in the human cell-type specific hypermethylated blocks (**Figure 3e - right**). Specifically, HOXB13 was the top TF associated with binding at sites within the human hypermethylated DMBs. Recently, HOXB13 has been found to control cell state through binding to super-enhancer regions, suggesting a novel regulatory function for cell-type specific hypermethylation(48). In addition to the common TFBS enriched by all cell-type specific blocks, endothelial-specific TFs were found to be enriched in the endothelial-cell hypomethylated blocks, including EWS, ERG, Fli1, ETV2/4, and SOX6 (see **Figure 4e**). Overall, these data indicate functions of these cell-type specific blocks that represent cell-specific biology that is still underexplored.

**Figure 4.**
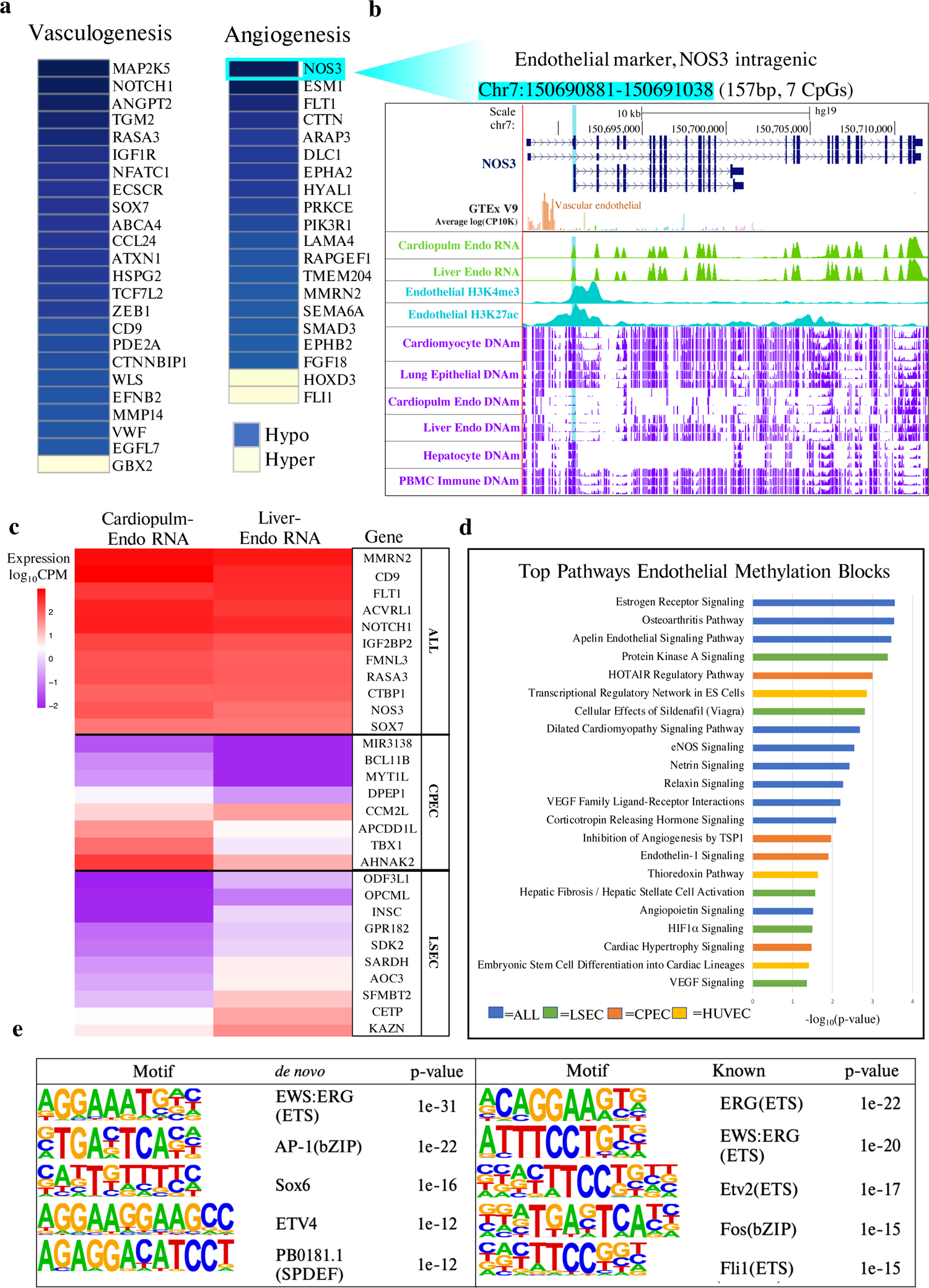
Methylation profiling of tissue-specific endothelial cell types reveals epigenetic heterogeneity associated with differential gene expression and biological functions. (a) Functions of genes adjacent to endothelial-specific methylation blocks (all p<0.05). Blue color indicates nearby hypomethylated regulatory blocks. Yellow color indicates nearby hypermethylated regulatory blocks. (b) Example of the NOS3 locus specifically unmethylated in endothelial cells. This endothelial-specific, differentially methylated block (DMB; highlighted in light blue) is 157 bp long (7 CpGs), and is located within the NOS3 gene, an endothelial-specific gene (upregulated in paired RNA-sequencing data as well as in vascular endothelial cells, GTEx inset). The alignment from the UCSC genome browser (top) provides the genomic locus organization and is aligned with the average methylation (purple tracks) across cardiomyocyte, lung epithelial, liver sinusoidal endothelial (LSEC), cardiopulmonary endothelial (CPEC), hepatocyte, and immune (PBMC) samples (n=3 / cell-type group). Results from RNA-sequencing generated from paired cell-types are depicted (green tracks) as well as peak intensity from H3K27ac and H3K4me3 published ChIP-seq data generated in endothelial cells (blue tracks). (c) Expression levels of genes adjacent to tissue-specific endothelial methylation blocks. Expression data were generated from paired RNA-sequencing of the same cardiopulmonary endothelial (CPEC) and liver sinusoidal endothelial (LSEC) cell populations used to generate methylation reference data. Pan-endothelial genes upregulated in both populations (ALL) are identified as common endothelial-specific methylation blocks to both LSEC and CPEC tissue-specific endothelial populations. (d) Pathways related to the biological function of genes containing endothelial-specific methylation blocks (all p<0.05). Unique pathways in distinct tissue-specific endothelial cells are highlighted in distinct colors. (e) Top 5 transcription factor binding sites enriched amongst endothelial-specific hypomethylated blocks, using HOMER de novo and known motif analysis. The background for the HOMER analysis consisted of 3,589 non-endothelial cell-type specific hypomethylated blocks. Abbreviations: CPEC = cardio-pulmonary endothelial cell, HUVEV = human umbilical vein endothelial cell, LSEC = liver sinusoidal endothelial cell.

### Methylation profiling of tissue-specific endothelial cell types reveals epigenetic heterogeneity associated with differential gene expression

Radiation-induced endothelial damage is thought to be a leading cause for development of late-onset complications from cardiovascular disease(49–51). We thus generated tissue-specific endothelial methylomes paired with transcriptomes to be able to identify damage to distinct populations of microvascular and large vessel endothelial cell types including coronary artery, pulmonary artery, cardiac microvascular, and liver sinusoidal endothelia. We also made use of publicly available liver sinusoidal endothelial(52) and umbilical vein endothelial methylomes(53) to complement our data (**Supplemental Table S1**). Previous studies support considering the heart and lung as an integrated system in the development of radiation damage due to the shared cardiopulmonary circulation(4). Therefore, we merged cardiac and pulmonary endothelial cell types to generate a joint cardiopulmonary endothelial signal and identified the specific methylation blocks for cardiopulmonary (CPEC), liver sinusoidal (LSEC), and umbilical vein endothelial (HUVEC) cell types as distinct populations. We also identified pan-endothelial methylation blocks with methylation status in common to all endothelial cell populations relative to other cell-types (**Supplemental Figure 6a-f**). Pathway analysis of genes associated with these genomic regions confirmed endothelial cell identity based on genes involved in the regulation of vasculogenesis and angiogenesis (**Figure 4a**). In addition, unique pathways identified the tissue-specific epigenetic diversity of endothelial cell populations from different organs (**Figure 4d**). The DNA methylation status at several tissue-specific blocks was found to correspond with RNA expression levels of known endothelial-specific genes, confirming the identity of endothelial populations characterized (**Figure 4b and 4c, Supplemental Table 9**)(46, 54–59). For example, hypomethylation was associated with increased expression at several pan-endothelial genes, including NOTCH1, ACVRL1, FLT1, MMRN2, NOS3 and SOX7. Likewise, hypomethylation at cardiopulmonary- or liver-specific endothelial genes was associated with differential expression when comparing the two populations to reflect tissue-specific differences.

### Methylated cfDNA changes indicate dose-dependent radiation damage in mice

To explore the relationship between radiation-induced damage in tissues to changing proportions of cell-free DNA origins in the circulation, we used mice to model the response to exposure from different radiation doses. Mice received upper thorax radiation of 3 Gy or 8 Gy relative to sham control, forming three groups for comparison (**Figure 1**). Tissues and serum were harvested 24 hours after the last fraction of treatment and tissues in the path of the radiation beam (heart, lungs and liver) were analyzed. H&E-stained sections showed a visible, dose-dependent impact of radiation on the tissues (**Figure 5a**). The changes were most apparent in tissue sections of the lungs showing noticeable alveolar collapse with increased radiation dose. Liver tissues showed increased fibrosis with increased radiation doses and only minor changes were apparent in cardiac tissues matching with its higher resilience to radiation. Tissue effects were also assessed through qPCR analysis of established indicators of radiation effects, including expression of CDKN1A (p21), that exhibited a dose-dependent increase in expression in response to radiation in all tissues (p<0.05, Kruskal-Wallis Test) (**Figure 5b, Supplemental Figure 4**)(60).

**Figure 5.**
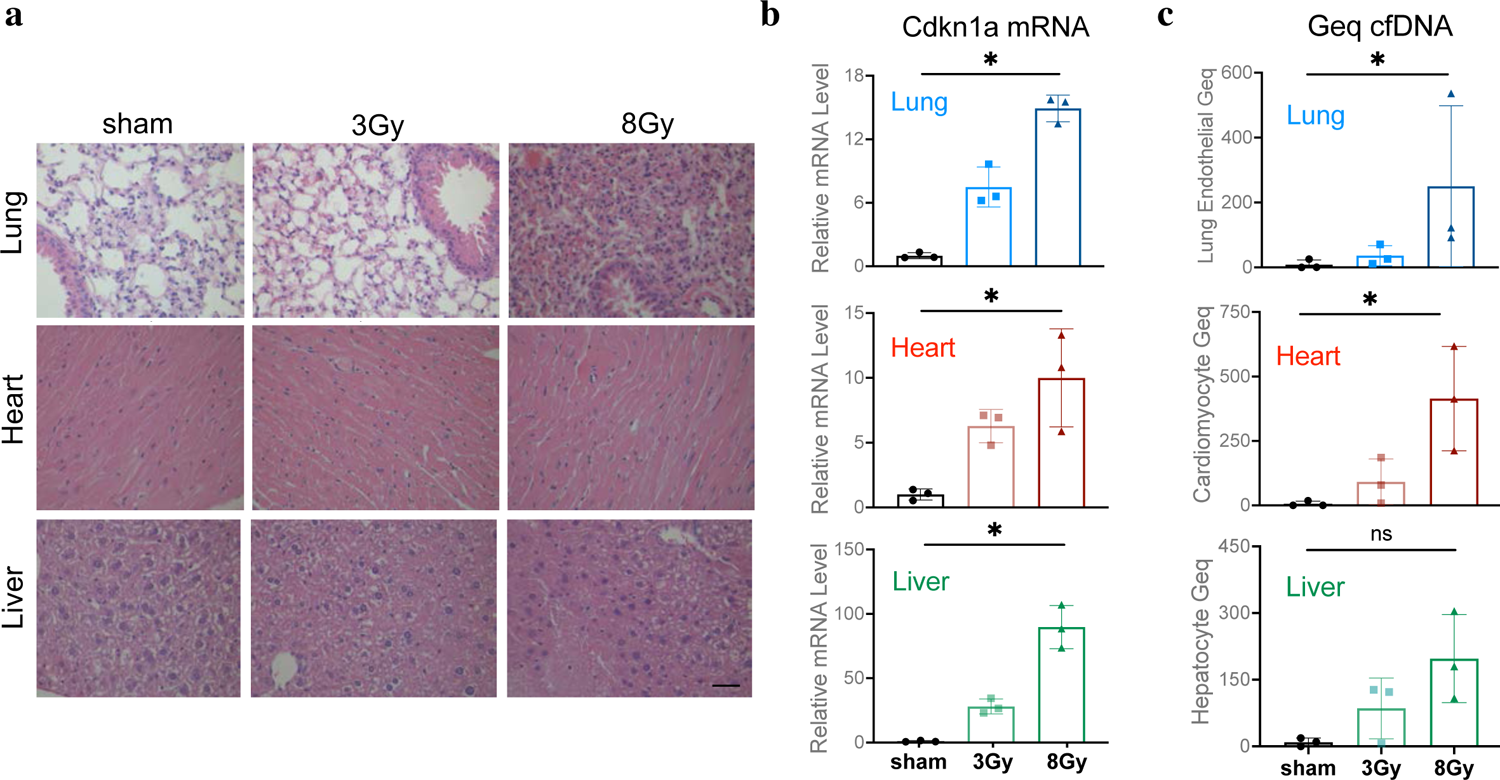
Dose-dependent radiation damage in mouse tissues correlates with the origins of methylated cfDNA in the circulation. (a) Representative H&E staining of lung, heart, and liver tissues from mice treated with 3Gy or 8Gy radiation compared to sham control. Scale bar, 200 μm. (b) qPCR analysis of CDKN1A (p21) mRNA. The expression in each sample was normalized to ACTB and is shown relative to the expression in the sham control. Mean ± SD; N = 3. Kruskal-Wallis test was used for comparisons amongst groups; lung tissue p=0.004, heart tissue p=0.025, liver tissue p=0.004. (c) Lung endothelial, cardiomyocyte and hepatocyte methylated cfDNA in the circulation of mice treated with 3Gy and 8Gy radiation compared to sham control expressed in Genome Equivalents per mL serum (Geq/mL). CfDNA was extracted from 18 mice (n=6 in each group) with cfDNA from 2 mice pooled in each methylome preparation. Mean ± SD; N = 3 independent methylome preparations. Kruskal-Wallis test was used for comparisons amongst groups. ns, P ≥0.05; *, P < 0.05; lung endothelial p=0.01, cardiomyocyte p=0.01, hepatocyte p=0.13.

To assess whether these damages of heart, lung, and liver are reflected in altered cfDNA patterns in the circulation, we used capture sequencing of CpG containing cfDNA fragments. For the data analysis we applied the above-described cell-type specific methylation blocks derived from the mouse methylation atlas. We found a significant dose-dependent increase in lung endothelial, cardiomyocyte and combined solid organ cfDNA across all three treatment groups that correlated with radiation-induced cell death in the corresponding tissues (p<0.05, Kruskal-Wallis Test) (**Figure 5c and Supplemental Figure 8g)**. The dose-dependent increase in hepatocyte cfDNA was not statistically significant and immune cell cfDNA showed no change between treatment groups (p≥0.05, Kruskal-Wallis Test) (**Figure 5c and Supplemental Figure 8f**). We conclude that changes in cfDNA fragments in the circulation can reveal the cellular source of radiation-induced damage in tissues.

### Radiation treatment of patients with breast cancer

To evaluate whether changes in cfDNA patterns could indicate damages to tissues in patients treated with radiation, we collected serum samples from randomly selected breast cancer patients at three timepoints during their standard-of-care radiation therapy after surgery **(Figure 1)**. A baseline sample was taken for each patient before onset of radiation therapy and a second End-Of-Treatment (EOT) sample was taken 30 minutes after the last treatment. Finally, a recovery sample was taken one month after completion of radiation therapy. Demographic information and clinical characteristics of patients enrolled in this study are in **Supplemental Table 8.**

### Methylated cfDNA changes provide an estimate of tissue dose to indicate radiation-induced damages to healthy tissues

Due to close proximity with the target treatment area, the heart and lungs are common organs-at-risk for breast cancer patients undergoing radiation therapy (**Figure 6a**). To assess therapy-induced lung damage, we examined cfDNA from serum for the presence of lung epithelial DMBs. Interestingly, we did not observe a significant increase in lung epithelial cfDNA across all patients (p≥0.05, Friedman Test) (**Figure 6b**). However, a few patients showed increased lung epithelial cfDNA indicating lung damage that correlated with increasing dose and volume of the lungs targeted (**Figure 6c**). Changes in lung epithelial cfDNA after radiation correlated with the volume of the ipsilateral lung receiving 20 Gy doses (Lung V20) (Pearson’s r = 0.78, p <0.05). In addition to lung injury, cardiovascular disease is one of the most serious complications from radiation exposure that is associated with increased morbidity and mortality(6). Through deconvolution using cardiopulmonary endothelial (CPEC) and cardiomyocyte-specific DMBs, we found increased CPEC and cardiomyocyte cfDNA in serum indicating significant cardiovascular cell damage across all breast cancer patients (p<0.05, Friedman Test) (**Figure 6d and f**). Changes in total endothelial cfDNA after radiation correlated with the average volume of the lung receiving a 5 Gy dose (Lung V5 Mean) (Pearson’s r = 0.71, p <0.05) (**Figure 6g**). Surprisingly, cardiomyocyte-specific methylated DNA in the circulation correlated with the maximum radiation dose to the heart (Pearson’s r = 0.63, p <0.05), but not the mean dose to the heart (Pearson’s r = −0.09, p≥0.05) (**Figure 6e**). This suggests that radiation-induced damage of cardiomyocytes requires a threshold dose indicating the relative resilience of this cell type compared to epithelial and endothelial cells from the heart and lungs.

**Figure 6.**
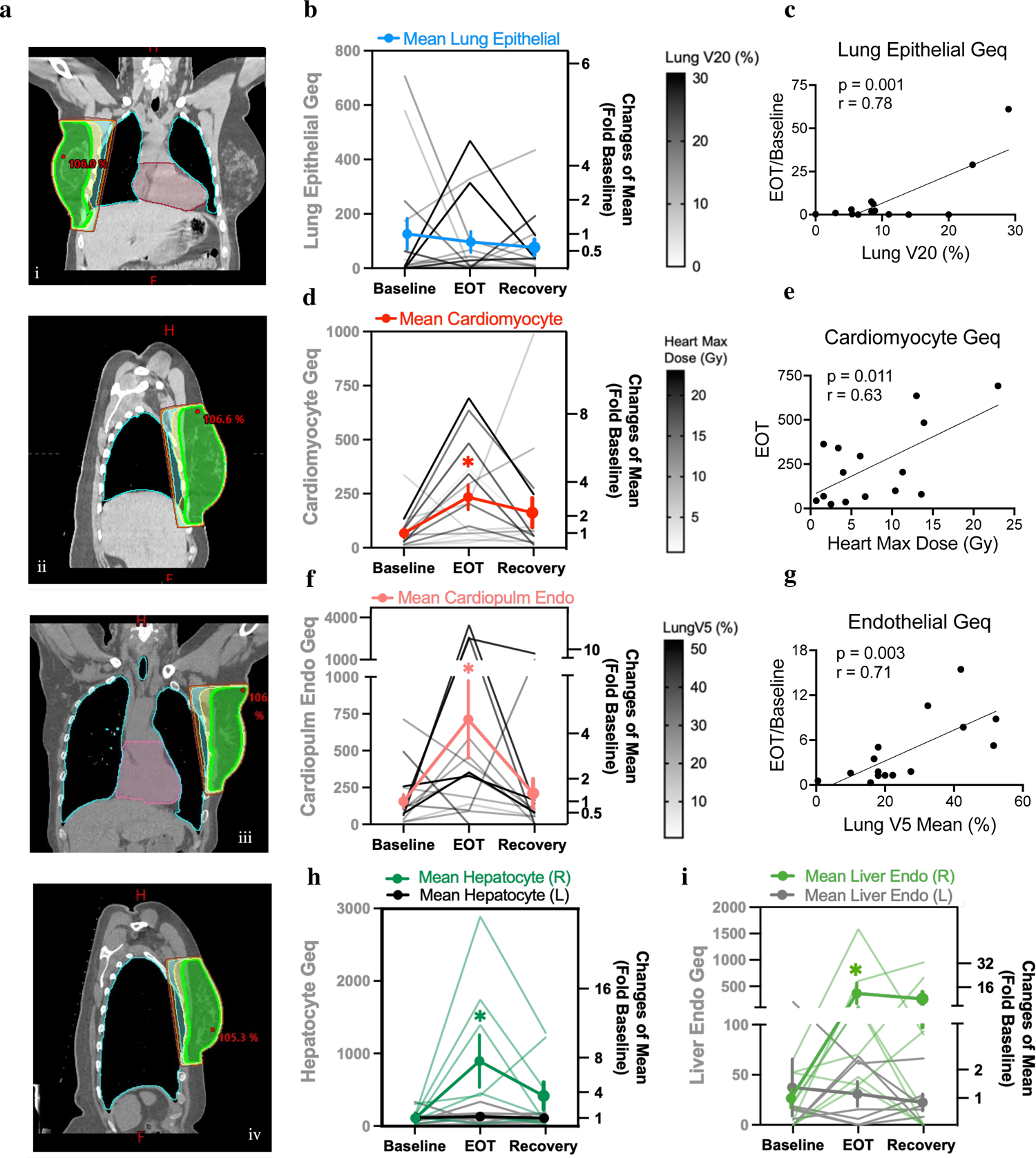
Methylated cell-type specific cfDNAs provide an estimate of tissue dose to indicate radiation-induced tissue damages in breast cancer patients. (a) Representative 3D-CRT treatment planning for patients with right-sided (i and ii) and left-sided (iii and iv) breast cancer, respectively. Computed tomography simulation of coronal and sagittal images depicting the anatomic position of the target volume in relation to nearby organs. The color map represents different radiation dose levels or isodose lines (Green: 95% of prescription dose, yellow: 90% isodose line, cyan: 80% isodose line, orange: 70% isodose line, brown: 50% isodose line). (b, d, f) Lung epithelial, cardiomyocyte, and cardiopulmonary endothelial (CPEC) cfDNA (in Geq/mL) in serum samples collected at different times. Fragment-level deconvolution used lung epithelial (n=69), cardiomyocyte (n=375), and CPEC specific methylation blocks (n=99), respectively. Friedman test was performed comparing paired results at baseline, EOT, and recovery timepoints. The results were considered significant when *p < 0.05; ns, p ≥ 0.05; lung epithelial p=0.99, cardiomyocyte p= 0.01, cardiopulmonary endothelial p=0.03. Mean fold change in lung epithelial, cardiomyocyte, and CPEC cfDNA at EOT and recovery relative to baseline levels is shown in bold. Data presented as mean ± SEM; N = 15. (b) Correlation of lung epithelial cfDNA with dosimetry data. EOT/Baseline represents the fraction of lung epithelial cfDNA post-radiation at end-of-treatment (EOT) relative to baseline levels. The volume of the lung receiving 20 Gy dose is represented by Lung V20 (%). One patient (RT108) had a five-fold increase in lung epithelial methylated DNA after radiation relative to all other patients and thus was treated as an outlier and removed from correlation analysis (with inclusion of RT108, Pearson’s r = 0.67 and p = 0.006; with exclusion of RT108, Pearson’s r = 0.78 and p = 0.001). (e) Correlation of cardiomyocyte cfDNA at EOT with the maximum dose to the heart (Gy). (g) Correlation of total endothelial cfDNA with dosimetry data as determined by deconvolution at pan-endothelial methylation blocks (n=131). EOT/Baseline represents the fraction of endothelial cfDNA post-radiation at end-of-treatment (EOT) relative to baseline levels. The mean volume of the lung receiving 5 Gy dose is represented by Lung V5 Mean (%). *(c, e, g) Pearson correlation r was calculated, and linear correlation was considered significant when *p < 0.05; ns, p ≥ 0.05. (h, i) Hepatocyte and Liver sinusoidal endothelial (LSEC) cfDNA (in Geq/mL) in serum samples collected at different times (EOT, end of treatment). Fragment-level deconvolution used top hepatocyte (n=200) and liver endothelial-specific methylation blocks (n=61). Mean fold change in right-sided hepatocyte and liver endothelial cfDNA at EOT and recovery relative to baseline levels is shown in bold. Data presented as mean ± SEM; N = 8 right-sided, N = 7 left-sided. Wilcoxon matched pairs signed rank test was used for comparison amongst groups and results were considered significant when *p < 0.05; ns, p ≥ 0.05; hepatocyte right-sided p= 0.02, hepatocyte left-sided p=0.81, LSEC right-sided p=0.02, and LSEC left-sided p=0.93.

### Radiation-induced hepatocyte and liver endothelial cfDNAs in patient with right-versus left-sided breast cancer

While liver damage is not a common radiation-induced toxicity in breast cancer patients, a substantial dose may still be administered to the liver, especially with right-sided tumors (**Figure 6a**). We used the top hepatocyte (n=200) and liver sinusoidal endothelial DNA methylation blocks to assess the sequence data for the presence of liver-derived cfDNA. Surprisingly, in patients receiving radiation treatment of right-sided breast cancer, an increase in hepatocyte plus liver sinusoidal endothelial methylated DNA in the circulation indicated significant radiation-induced cellular damage in the liver (p<0.05, Wilcoxon matched-pairs signed rank test) (**Figure 6h and i**). Elevated levels of either hepatocyte and/or liver sinusoidal endothelial cfDNA were detected in seven of the eight breast cancer patients with right-sided tumors. In contrast, there was no significant increase in hepatocyte or liver sinusoidal endothelial cfDNA in patients with left-sided breast cancer (p≥0.05, Wilcoxon matched-pairs signed rank test).

### Distinct endothelial and epithelial damages from radiation

We observed distinct epithelial and endothelial cell-type responses to radiation across the different tissue cfDNAs profiled. Different responses to radiation were observed when comparing hepatocyte to lung epithelial damages (**Fig 6b versus 6h)**, demonstrating the ability of methylated cfDNA to distinguish between tissue-specific epithelial cell types from serum samples. Likewise, analysis for tissue-specific endothelial populations revealed differences in cardiopulmonary microvascular and liver sinusoidal endothelial responses to radiation (**Fig 6f versus 6i**). In general, there was greater magnitude of damage to the combined endothelium compared to the epithelium across different organs (**Supplemental Figure 8e**). The endothelium forms a layer of cells lining blood as well as lymphatic vessels and turnover of this cell type may contribute to the high amplitude of signal detected from serum(24). This could, however, also be a result of the different sensitivities of endothelial versus epithelial cell types to radiation-induced damage. There was a five-fold higher signal from cardiopulmonary endothelial cfDNA compared to lung epithelial cfDNA. Also, in comparison to lung epithelial- and cardiopulmonary endothelial-derived cfDNA, sustained injury and delayed recovery is indicated by elevated cardiomyocyte cfDNA at the recovery time point (two-fold elevation from baseline) (**Figure 6d)**. This may reflect important differences in cell turnover rates leading to differential processes of regeneration and repair in these cell types. Notably, one month after completion of radiation therapy, lung epithelial and cardiopulmonary endothelial damage signatures detected from cfDNA had returned to baseline levels whereas sustained higher cfDNA from cardiomyocytes and liver cell-types indicates lingering tissue remodeling. Taken as a whole, these findings demonstrate applicability of this approach to uncover distinct cellular damages in different tissues during the course of treatment by the analysis of blood samples.

## Discussion

This study demonstrates the feasibility of tissue-of-origin analysis of cell-free methylated DNA to monitor tissue responses to radiation exposure. The assignment of DNA fragments extracted from serum samples from patients undergoing treatment as well as from experimental animals to specific cell types required in-depth analysis of tissue- and cell-type DNA methylation patterns. We expected to at least identify a set of biomarkers from this analysis and were positively surprised by the significant association of the cell-type specific DNA methylation blocks with cell-type specific gene expression, transcription factor binding motifs and signaling pathway regulation. We were particularly intrigued by how well-conserved cell-type specific DNA methylation appears to be across different individuals suggesting broad applicability in the monitoring of tissue damage in diverse patient populations. It appears that disease- or ageing-related changes in DNA methylation occur outside the cell-type specific blocks and thus will exert their impact without altering the features defining a particular cell type.

To enhance the quality and sensitivity of the analyses, we developed human and mouse reference methylation atlases that were tailored to this application and study design. We optimized a capture-sequencing methodology for low-integrity cfDNA samples from human and mouse serum samples, achieving increased sequencing coverage and sensitivity of deconvolution applying a fragment-level probabilistic model. These improvements allow for accurate cellular assignment of cfDNA fragments in serum. When evaluating our approach, we also directly compared cfDNA extracted from serum and plasma samples harvested from the same donor. The results were highly correlated and we found slightly less variation across donors in the cell-type proportions contributing to the cfDNA extracted from serum compared to plasma (details in the Supplemental Methods section; **Supplemental Figure 9**). Overall, the sequencing-based fragment-level deconvolution model allows for accurate prediction of contributions from solid organ cell types.

Comparing the origins of elevated cfDNA after radiation showed similar changes in both human and mouse serum samples, providing further validation. In both human and mouse, there was a significant increase in endothelial and cardiomyocyte cfDNA after radiation. Likewise, there was an overall increase in cfDNA derived from any solid-organ tissue post-radiation (**Supplemental Figure 8**). We also detected significantly increased breast basal and luminal epithelial cfDNA across all breast cancer patients (**Supplemental Figure 8c and d**). While there wasn’t a significant dose-dependent increase, there was a parallel increase in mammary epithelial cfDNA in mice treated with radiation (3Gy and 8Gy combined) compared to sham control (**Supplemental Figure 8h**). The total concentration of cfDNA was elevated in some breast cancer patients at the end of treatment (EOT) as well, suggesting an overall increase in cfDNA shortly after radiation treatment in some patients (**Supplemental Table 7**). However, this trend was not consistent for all patients. Similarly, changes in mouse cfDNA concentration with increasing radiation dose were not significant (p≥ 0.05, Kruskal-Wallis Test; **Supplemental Table 6**) as also found by others(61). However, despite similarities, it was difficult to fully align the human and mouse results given the scarcity of deep-sequencing WGBS data from purified murine cell types.

We generated tissue-specific endothelial methylomes to profile differing sensitivities of cardiopulmonary microvascular and liver sinusoidal endothelial cells to radiation. In many tissues, the vascular endothelium is among the first cell types known to be damaged(4, 62). Likewise, our analysis indicated greater damage to the endothelium compared to epithelium in different organs. However, the role of radiation-induced endothelial damage in mediating acute and chronic adverse effects has yet to be fully understood. While there are important tissue-specific differences between these endothelial cell populations reflected by several distinct methylation patterns identified in our atlas, we were unsure if we would be able to discriminate contributions between these cell-types in the circulation given the high similarity of methylation patterns across all endothelial cell-types derived from common lineages (**Figure 2a, b**). We were pleasantly surprised by the diverse injury patterns observed juxtaposing cardiopulmonary endothelial to liver sinusoidal endothelial signals that were consistent with our expectations based on corresponding clinical data. Recent studies suggest that compromised endothelial cell function also impairs wound healing by depriving tissues of signals necessary for regeneration and contributing to accelerated aging of the hematopoietic and vascular systems(63–65). Notably, one month after completion of radiation therapy we found that the majority of damage signatures in the circulation had returned to baseline levels whereas increased turnover of cardiomyocytes and liver cell-types is indicated by sustained elevation of the respective cfDNAs likely due to ongoing tissue remodeling. Exploration of these tissue- and cell-type specific differences may shed light on previously unknown mechanisms of radiation-induced damages.

The liver is not a common organ-at-risk for radiation-induced toxicity in breast cancer, and we were surprised to find an increase in hepatocyte and liver sinusoidal endothelial cell methylated DNA in the circulation of patients receiving radiation treatment for right-sided breast cancer. Previous studies have not found a meaningful relationship between breast cancer RT and overt liver fibrosis, even at doses higher than 40 Gy; although, increased hepatic exposure is expected in radiation treatment of patients with right-sided breast cancer(66). In comparison to previous methods that did not detect radiation-induced liver damage, our finding demonstrates the sensitivity of circulating cell-free methylated DNA to identify previously unknown cell types and tissues impacted by radiation treatment. Despite being at subclinical levels, this may become relevant in patients with hepatotoxic therapy regimens or co-morbidities.

While the methylated cfDNA signals indicate damage to liver cells associated with treatment of right-sided tumors, it is unknown if this contributes to clinically impactful adverse effects. Interestingly, three patients (RT-102, RT-103, RT-107) presented with grade 2 skin toxicity based on Common Terminology Criteria for Adverse Events (CTCAE V5) within our study timeline up to one month after completion of radiation therapy. We detected significantly increased breast basal and luminal epithelial damages in these three patients to correlate with the clinical presentation (**Supplemental Figure 8c and d**). We also detected elevated breast epithelial injury in patients that underwent mastectomy, were treated with proton beam therapy, and had higher overall dose administered – all clinical factors associated with elevated risk of skin toxicity. With adaptive radiation planning, the potential to modify ongoing treatment could tailor therapy to individual needs, both increasing dose in more tolerant patients and decreasing dose in sensitive patients between fractions to optimize efficacy while also minimizing toxicity. There are many factors to consider when assessing the risk for adverse events, including patient characteristics, pre-existing disease, and treatment-related risk factors, such as dose, tumor location and effects of combination therapy. Cell-free DNA analyses may be used to compare these different cohorts as well as distinct toxicity profiles associated with 3D-CRT to other more recent RT techniques, such as proton beam therapy (PBT) and intensity-modulated radiation therapy (IMRT). Likewise, exploration of regional variation in tissue-specific response to radiation may offer new opportunities to reduce normal tissue side effects(67).

As proof of concept, radiation treatment served as a powerful tool to validate the cell-type specific methylation signatures developed in the atlases and demonstrate application to detect cellular injury in the circulation. In mice, we performed paired tissue and serum analysis allowing for a direct comparison. Similarly, treatment planning for breast cancer patients provided an estimate of the organ volume impacted and radiation dose level for organs-at-risk from radiation damage, including the heart and lungs. Given this tissue-informed knowledge, we observed a striking degree of correlation between dose and cell-type contributors to the circulation after radiation, supporting that methylated cfDNA changes can indicate the actual tissue dose administered providing an objective measure of cellular injury in vivo. We conclude that the minimally invasive detection of cell-free methylated DNA from serum samples can indicate organ-specific damages, biologically effective radiation doses received by tissues and reveal previously unknown cell types impacted by radiation treatment.

## Methods

### Human BC patient serum sample collection

Serial serum samples were collected from 15 breast cancer patients at Baseline (before radiation treatment), End-of-Treatment (EOT; 30 minutes after the last radiation treatment), and Recovery (one month after cessation of radiation treatment), thus allowing for a within-patient internal control and baseline. A schematic of the time series for sample collection can be found in **Figure 1**. For serum isolation, peripheral blood (∼8-12 mL) was collected in red-top venous puncture tubes and allowed to clot at room temperature for 30 minutes before centrifugation at 1500 x g for 20 min at 4°C to separate the serum fraction. Patients received either three-dimensional conformal RT (3D-CRT) or a combination of proton beam therapy (PBT) and 3D-CRT. Patient characteristics and treatment details including radiation dosimetry are summarized in **Supplemental Tables 8 and 12.**

### Mouse serum and tissue collection

C57Bl6 mice (n=18) were irradiated to the upper thorax at different doses (sham control, 3Gy, 8Gy) for 3 consecutive treatments. Serum and tissues were collected 24 hours after the last radiation dose. For serum isolation, blood was collected via cardiac puncture (∼1mL) and allowed to clot at room temperature for 30 minutes before centrifugation at 1500 x g for 20 min at 4°C to separate the serum fraction. Heart, lung, and liver tissues were harvested and sectioned to be both flash frozen and formalin fixed for subsequent analysis.

### Cell isolation to generate reference methylomes

Reference methylomes were generated for mouse immune cell types and human endothelial cell types to complement publicly available datasets. Peripheral blood and bone marrow were isolated and spleens from healthy C57Bl6 mice were dissociated to single cells and FACS sorted using cell-type specific antibodies. Buffy coat (n=4), bone marrow (n=3), CD19+ B cell (n=1), CD4 T cell (n=1), CD8 T cell (n=1) and Gr1+ Neutrophil (n=1) methylomes were generated after cell isolation using the following antibodies: FITC anti-mouse CD45 (cat#103107), Alexa Fluor 647 anti-mouse CD3 (cat#100209), Brilliant Violet 711 anti-mouse CD4 (cat#100549), Brilliant Violet 421 anti-mouse CD8a (cat#100737), PE anti-mouse CD19 (cat#152407), PE/Cy7 anti-mouse Ly-6G/Ly-6C (Gr-1) (cat#108415) (all BioLegend 1:20). Cryopreserved passage 1 human liver sinusoidal endothelial cells (LSEC) were purchased from ScienCell research laboratories (SKU#5000). Purity was determined by immunofluorescence with antibodies specific to vWF/Factor VIII and CD31(PECAM). Cryopreserved passage 2 human coronary artery (HCAEC SKU#C-14022), cardiac microvascular (HCMEC SKU#C-14029), and pulmonary artery endothelial cells (HPAEC SKU#C-14024) were isolated from single donor healthy human tissues purchased from PromoCell. Paired RNA-seq data was generated from the same cell-populations used for DNA methylation profiling to validate the identity of purchased cell populations through analysis of cell type expression markers.

### Isolation of circulating cell-free DNA (cfDNA)

Circulating cell-free DNA was extracted from 3 to 4 mL human serum or plasma or 0.5 mL mouse serum, using the QIAamp Circulating Nucleic Acid kit (Qiagen) according to the manufacturer’s instructions. Cell-free DNA was quantified via Qubit fluorometer using the dsDNA High Sensitivity Assay Kit (Thermo Fisher Scientific). Fragment size distribution of isolated cfDNA was validated on the 2100 Bioanalyzer TapeStation (Agilent Technologies). Additional size selection using Beckman Coulter beads was applied to remove high-molecular weight DNA reflective of cell-lysis and leukocyte contamination as previously described(33). Size distribution of cell-free DNA fragments was re-verified following purification (**Supplemental Figure 9**). Additional details can be found in Supplemental Methods.

### Isolation and fragmentation of genomic DNA

Genomic DNA from tissues was extracted with the DNeasy Blood and Tissue Kit (Qiagen) following the manufacturer’s instructions and quantified via the Qubit fluorometer dsDNA BR Assay Kit (Thermo Fisher Scientific). Genomic DNA was fragmented via sonification using a Covaris M220 instrument to the recommended 150-200 base pairs before library preparation. Lambda phage DNA (Promega Corporation) was also fragmented and included as a spike-in to all DNA samples at 0.5%w/w, serving as an internal unmethylated control. Bisulfite conversion efficiency was calculated through assessing the number of unconverted C’s on unmethylated lambda phage DNA. The SeqCap Epi capture pool contains probes to capture the lambda genomic region from base 4500 to 6500.

### Bisulfite capture-sequencing library preparation

Bisulfite capture-sequencing libraries were generated from either cfDNA or fragmented genomic DNA using the same protocol. As a first step, WGBS libraries were generated using the Pico Methyl-Seq Library Prep Kit (Zymo Research) with the following modifications: Bisulfite-conversion was carried out using the EZ DNA Methylation Gold kit (Zymo Research) instead of the EZ DNA Methylation-Lightning Kit. For mouse samples, cfDNA from two mice in the same group was pooled as the input of library preparation. An additional 2 PCR cycles were added to the recommended cycle number based on the total amounts of input cfDNA. WGBS libraries were eluted in 15uL 10mM Tris-HCl buffer, pH 8. Library quality control was performed with an Agilent 2100 Bioanalyzer and quantity determined via the KAPA Library Quantification Kit (KAPA Biosystems).

Cell-free WGBS libraries were pooled to meet the required 1ug DNA input necessary for targeted enrichment. However, no more than four WGBS libraries were pooled in a single hybridization reaction and the 1ug input DNA was divided evenly between the libraries to be multiplexed. Hybridization capture was carried out according to the SeqCap Epi Enrichment System protocol (Roche NimbleGen, Inc.) using SeqCap Epi CpGiant probe pools for human samples and SeqCap Epi Developer probes for mouse samples with xGen Universal Blocker-TS Mix (Integrated DNA Technologies, USA) as the blocking reagent. Washing and recovering of the captured library, as well as PCR amplification and final purification, were carried out as recommended by the manufacturer. The capture library products were assessed by Agilent Bioanalyzer DNA 1000 assays (Agilent Technologies, Inc.). Bisulfite capture-sequencing libraries with inclusion of 15-20% spike-in PhiX Control v3 library (Illumina) were clustered on an Illumina Novaseq 6000 S4 flow cell followed by 150bp paired-end sequencing.

### Bisulfite sequencing data alignment and preprocessing

Paired-end FASTQ files were trimmed using TrimGalore (V 0.6.6) with parameters “-- paired -q 20 --clip_R1 10 --clip_R2 10 --three_prime_clip_R1 10 --three_prime_clip_R2 10” (https://github.com/FelixKrueger/TrimGalore). Trimmed paired-end FASTQ reads were mapped to the human genome (GRCh37/hg19 build) using Bismark (V 0.22.3)(34) with parameters “--non-directional”, then converted to BAM files using Samtools (V 1.12)(35). BAM files were sorted and indexed using Samtools (V1.12). Reads were stripped from non-CpG nucleotides and converted to BETA and PAT files using *wgbstools* (V 0.1.0), a tool suite for working with WGBS data while preserving read-specific intrinsic dependencies (https://github.com/nloyfer/wgbs_tools)(22, 36).

### Reference DNA methylation data from healthy tissues and cells

Controlled access to reference WGBS data from normal human tissues and cell types was requested from public consortia participating in the International Human Epigenome Consortium (IHEC)(37) and upon approval downloaded from the European Genome-Phenome Archive (EGA), Japanese Genotype-phenotype Archive (JGA), database of Genotypes and Phenotypes (dbGAP), and ENCODE portal data repositories. Reference mouse WGBS data from normal tissues and cells were downloaded from selected GEO and SRA datasets. Additional information and citation of reference methylation data used in this study can be found in Supplemental Methods and **Supplemental Tables 1 and 2**.

### Segmentation and clustering analysis

We segmented the genome into blocks of homogenous methylation as described by Loyfer et al 2022 using *wgbstools* (with parameters segment --max_bp 5000)(22, 36). In brief, a multi-channel Dynamic Programming segmentation algorithm was used to divide the genome into continuous genomic regions (blocks) showing homogenous methylation levels across multiple CpGs for each sample. We applied the segmentation algorithm to 297 human reference WGBS methylomes and retained 351,395 blocks covered by the hybridization capture panel used in the analysis of cfDNA that captures 80Mb (∼20% of CpGs). Likewise, segmentation of 109 mouse WGBS datasets from healthy cell types and tissues identified 1,344,889 blocks covered by the mouse hybridization capture panel that captures 210 Mb (∼75% of CpGs). The human blocks had a median length of 326 bp (interquartile range, IQR = 890 bp) and 8 CpGs (IQR = 14 CpGs). Similarly, the mouse blocks had a median length of 770 bp (IQR =1,252 bp) and 7 CpGs (IQR = 7 CpGs). The hierarchical relationship between reference tissue and cell type WGBS datasets was visualized as a tree dendrogram. The top 30,000 most variable methylation blocks containing at least three CpG sites and coverage across 90% of samples were selected. We computed the average methylation for each block and sample using wgbstools (-- beta_to_table). Trees were assembled using the unweighted pair-group method with arithmetic mean (UPGMA)(38), using scipy (V 1.7.1)(39) and L1 distance, and then visualized in R with the ggtree package (V 2.4.1)(40). The similarity between samples was assessed by the degree of variation in distance between samples of the same cell-type (average 23,056) compared to samples between different cell-types (average 273,018). Dimensional reduction was also performed on the selected blocks using the UMAP package (V 0.2.8.2.0). Default UMAP parameters were used (15 neighbors, 2 components, Euclidean metric, and a minimum distance of 0.1).

### Identification of cell-type specific methylation blocks

Tissue and cell-type specific methylation blocks were identified from reference WGBS data using custom scripts (**Supplemental code and Supplemental Methods**). We performed a one-vs-all comparison to identify differentially methylated blocks unique for each group. This was done separately for human and mouse. From this we first identified blocks covering a minimum of three CpG sites, with length less than 2Kb and at least 10 observations. Then, we calculated the average methylation per block/sample, as the ratio of methylated CpG observations across all sequenced reads from that block. Differential blocks were sorted by the margin of separation, termed “delta beta”, defined as the minimal difference between the average methylation in any sample from the target group vs all other samples. We selected blocks with a delta-beta ≥ 0.4 for human and ≥ 0.35 for mouse. Additional separation of endothelial cell populations from different tissues was performed to identify unique markers for liver endothelial versus cardiopulmonary endothelial blocks that do not overlap. Separately, pan-endothelial blocks were identified with methylation status in common to all endothelial cell populations. For some cell-types, a reduced subset of blocks (ie. top 200) were used for deconvolution in the circulation if the original number identified was greater than one standard deviation above the mean. Selected human and mouse blocks for cell types of interest can be found in **Supplemental Tables 3 and 4.**

### Likelihood-based probabilistic model for fragment-level deconvolution

The cell type origins of cfDNA were determined using a probabilistic fragment-level deconvolution algorithm. Using this model, the likelihood of each cfDNA molecule was calculated using a 4^th^-order Markov Model, by which methylation of each CpG site directly depends on up to four adjacent previous sites within each fragment. We estimated these parameters for each differential block at every tissue and cell-type, and then used Bayes’ theorem to infer the posterior probability of cell-of-origin for each fragment, based on its complete methylation pattern. The model was trained on reference bisulfite-sequencing data from normal cells and tissues of known identity to learn the distribution of each marker in the target tissue/cell population of interest compared to background. Then the model was applied to predict the origins of each cfDNA molecule. The joint probability of each cfDNA molecule (methylation patterns and cellular origin) is calculated based on the likelihood of the methylation pattern (using the parameters for that cell type) times the prior probability that a read is originating from the target cell type. A prior of 0.1 was used for the combined endothelial cell-type group and 0.85 for the combined immune cell-type group as expected based on findings in previous reports(24). A prior of 0.05 was used for all other solid organ cell-types. Finally, each fragment is assigned to the cell type of origin with the maximal posterior probability (“hard” assignment). The proportion of molecules (fragments) assigned to the tissue of interest across all cell-type specific markers was then averaged and used to determine the relative abundance of cell-free DNA derived from that tissue in each respective sample. We then adjusted the resulting proportions from all cell types to have a sum of 1 by imposing a normalization constraint. Tissue-specific endothelial cell-types were normalized within the predicted total endothelial proportion identified by pan-endothelial markers in common to all endothelial cell-types. Predicted cell-type proportions were converted to genome equivalents and reported as Geq/mL through multiplying the concentration of cfDNA (ng/mL) by the mass of the human haploid genome 3.3 x 10^-12^ grams or the mouse haploid genome equivalent of 3.0 x 10^-12^ grams.

### In-silico simulations and WGBS deconvolution

In silico mix-in simulations were performed using *wgbstools* (V 0.1.0)(36) to validate the fragment-level deconvolution algorithm at the identified cell-type specific blocks (**Supplemental Figures 3, 5, 6 and 7**) as previously described(22, 24). For each cell type profiled, we mixed known proportions of target fragments into a background of leukocyte fragments using wgbstools mix_pat. The leukocyte fragments were obtained from n=4 buffy coat samples in mouse and n=5 buffy coat samples in human. We performed three replicates for each admixture ratio assessed (0.05%, 0.1%, 0.5%, 1%, 2%, 5%, 10%, 15%), which were analyzed as described above, and present the average predicted proportion and standard deviation across all replicates. Model accuracy was assessed through correct classification of the actual percent target mixed.

### Functional annotation and pathway analysis

Cell-type specific methylation blocks were annotated and motif analysis was performed using HOMER (V4.11.1) (http://homer.ucsd.edu/homer/)(41) using the annotatePeaks.pl and findMotifsGenome.pl functions. The top 5 motifs based on p-value were selected from each analysis. Pathway analysis of genes adjacent to identified tissue and cell-type specific methylation blocks was performed using Ingenuity Pathway Analysis (IPA)(42) (Qiagen) and Genomic Regions Enrichment of Annotations Tool (GREAT)(43). GeneSetCluster was used to cluster identified gene-set pathways based on shared genes(44).

### Genome Browser visualization

Endothelial methylomes and paired transcriptomes were uploaded as custom tracks for visualization on the UCSC genome browser(45). Methylomes were converted to bigwig format using the wgbstools beta2bw. Enrichment for chromatin marks was assessed through analysis of published H3K27ac and H3K4me3 ChIP-seq data(46). GTEx single nucleus RNAseq data was acquired from the GTEx v9 Portal (gtexportal.org)(47) and analyzed using R (V 4.1.3). Counts per ten thousand reads (CP10K) of NOS3 were log-transformed and averaged for each specific cell type. Color represents the general cell type and intensity of color represents the number of cells expressing NOS3.

#### Statistics

Statistical analyses for group comparisons and correlations were performed using Prism (GraphPad Software, Inc., United States) and R (V 4.1.3). A correlation analysis was performed to assess relationship between changing cell-free methylated DNAs and dose using Pearson’s Correlation Coefficient. Statistically significant comparisons are shown, with significance defined as P less than 0.05.

#### Study Approval

Breast cancer patients undergoing adjuvant radiation therapy were enrolled and provided signed informed consent in this IRB approved study at Medstar Georgetown University Hospital (IRB protocol # 2013-0049). Animal studies were approved by the Georgetown University IACUC (protocol #2017-0029).

## Author Contributions

MEB, AJK, SMR, EB, KU, and SR facilitated collection and processing of enrolled breast cancer patient samples. EB performed chart review and organized breast cancer patient data. MEB, AJK, and SMR performed bisulfite-capture sequencing library preparation. MEB and AJK purified cell-types and performed reference methylation data generation and validation. YL and HL facilitated radiation-treatment of mice. MEB and MOS facilitated serum and tissue collection from mice. AJK performed RT-qPCR and RNA-sequencing analysis. NS performed histological analysis of mouse tissues. MEB and NL performed bioinformatics analysis of WGBS data. MEB processed data for identification of mouse and human cell-type specific differential methylation. NL and TK created *wgbstools* for bioinformatics read-specific processing of WGBS data. NL and TK generated all scripts for WGBS data processing, segmentation, and identification of cell-type specific methylation blocks. NL, SSP, and TK developed the fragment-level deconvolution model for cfDNA analysis. SJ, APM, and ADM advised on statistical and computational data analysis. MEB and AW wrote the manuscript and generated all figures. AW, AR and KU conceived the study design and provided interpretation of results. Relative overall contribution was used as the method to assign the authorship order. All authors critically reviewed and approved the manuscript.

## Declarations

### Availability of data and materials

All data and code will be made available upon publication of this manuscript.

## Acknowledgements

Valuable discussions and suggestions were contributed by Habtom Ressom, Michael R. Lindberg, and Garrett T. Graham (all at Georgetown University). The graphical abstract and sample collection schema in this manuscript were created with BioRender.com.

We also acknowledge the following consortia that generated data used in this study: *KNIH -* This study makes use of data generated by the Korea Epigenome Project. A full list of the investigators who contributed to the generation of the data is available from 152.99.75.168/KEP. Funding for the project was provided by KOREA EPIGENOME PROJECT.

*AMED-CREST / NBDC -* A part of the data used for this research was originally obtained by AMED-CREST International Human Epigenome Consortium (IHEC) research project/group led by Prof. /Dr. Yae Kanai and by Core Research and Evolutional Science and Technology (CREST), International Human Epigenome Consortium (IHEC) project/group led by Hiroyuki Sasaki, both available at the website of the National Bioscience Database Center (NBDC; http://biosciencedbc.jp/en/) of the Japan Science and Technology Agency (JST).

*CUHK CNARG -* This study makes use of data generated by The Chinese University of Hong Kong (CUHK) Circulating Nucleic Acids Research Group, as reported by Cheng THT et al in Clin Chem (Cheng THT et al. Clin Chem. 2019 Jul;65(7):927-936)” and by Chen THT et al in Clin Biochem. 2017;50:496–501”.

*ENCODE -* We downloaded the call sets form the ENCODE portal (https://www.encodeproject.org/) with the following identifiers: ENCSR656TQD, ENCSR328TBS, ENCSR116JEF, ENCSR116JEF, ENCSR999RWT, ENCSR577RUU, ENCSR267SNS, ENCSR899RXH, ENCSR258MDR, ENCSR641SDF, ENCSR835OJU, ENCSR442FIY, ENCSR893RHD, ENCSR784VGW, ENCSR211VXF, ENCSR749COU, ENCSR733ZTZ, ENCSR728AGC.

*CEEHRC -* The results published here are in part based upon data generated by The Canadian Epigenetics, Epigenomics, Environment and Health Research Consortium (CEEHRC) initiative funded by the Canadian Institutes of Health Research (CIHR), Genome BC, and Genome Quebec. Information about CEEHRC and the participating investigators and institutions can be found at http://www.cihr-irsc.gc.ca/e/43734.html.

*DEEP -* This study makes use of data generated by the Deutsches Epigenom Programm DEEP. A full list of the investigators who contributed to the generation of the data is available from the consortium website (www.deutsches-epigenom-programm.de).

*Blueprint -* This study makes use of data generated by the Blueprint Consortium. A full list of the investigators who contributed to the generation of the data is available from www.blueprint-epigenome.eu. Funding for the project was provided by the European Union’s Seventh Framework Programme (FP7/2007-2013) under grant agreement no 282510 – BLUEPRINT.

## Supplementary Methods and Figures

Supplement to: “Cell-free, methylated DNA in blood samples reveals tissue-specific cellular damage from radiation treatment.” Barefoot *et al.* 2022.

### Supplemental Materials and Methods

#### Processing of human serum and plasma samples

Circulating cell-free DNA was extracted from 3 to 4 mL human serum or plasma or 0.5 mL mouse serum, using the QIAamp Circulating Nucleic Acid kit (Qiagen) according to the manufacturer’s instructions. Cell-free DNA was quantified via Qubit fluorometer using the dsDNA High Sensitivity Assay Kit (Thermo Fisher Scientific). Fragment size distribution of isolated cfDNA was validated on the 2100 Bioanalyzer TapeStation (Agilent Technologies). Additional size selection using Beckman Coulter beads was applied to remove high-molecular weight DNA reflective of cell-lysis and leukocyte contamination as previously described(1). Size distribution of cell-free DNA fragments was re-verified following purification.

Control human serum and plasma from healthy adult donors was purchased from Innovative Research (SKU#ISERS10ML and SKU#IPLASK2E10ML) to compare results from our analyses across sample preparations. While plasma is produced when whole blood is collected in tubes that are treated with anticoagulant, serum is obtained after allowing blood to clot for 30 minutes at room temperature and then centrifuging the samples to remove the cellular component(2, 3). Studies demonstrate that cellular components significantly increase in samples that sit longer than 60 minutes while clotting; however, adherence to standard operating procedures for preparation of serum and plasma have been found to greatly reduce contamination and sources of error(4). We took extra steps to address these concerns by ensuring timely processing of blood samples and performing an additional bead purification after cfDNA isolation to remove high-molecular weight DNA, likely derived from contaminating blood cell lysis (**Supplemental** Figure 9a and b). We found that taking this approach, cfDNA methylation status at the block level is highly correlated when comparing cfDNA derived from serum or plasma (**Supplemental** Figure 9c; Pearson r = 0.95). In addition, deconvolution analysis verified that the %immune and %solid organ origins of cfDNA does not vary across the two sample types (**Supplemental** Figure 9d). In fact, there appears to be slightly less variation across donors in the predicted cell type proportions composing cfDNA extracted from serum compared to plasma. Thus, despite an overall higher Geq Immune found in serum due to the overall higher cfDNA concentrations, this background signal is consistent from sample-to-sample allowing for accurate comparison of changes over time in serial samples collected from the same individuals (**Supplemental** Figure 9d and e).

#### RNA isolation, RNA-sequencing, and RT-qPCR analysis

RNA was isolated from tissues or sorted cells using the RNeasy Kit (Qiagen) following homogenization using the MagNA Lyser (Roche) according to the manufacturer’s protocol and quantified by Qubit RNA BR assay (Thermo Fisher Scientific). Total RNA was validated using an Agilent RNA 6000 nano assay on the 2100 Bioanalyzer TapeStation (Agilent Technologies). The resulting RNA Integrity number (RIN) of samples selected for downstream qPCR or RNAseq analysis was at least 7. Reverse transcription (RT) was done using the iScript cDNA Synthesis Kit (Bio-Rad) according to the manufacturer’s protocol. Real-time quantitative RT–PCR was performed with iQ SYBR Green Supermix (Bio-Rad). Primers used for RT–qPCR were purchased from Integrated DNA Technologies, and their sequences are provided below. Fold change was calculated as a percentage normalized to housekeeping gene human actin (ACTB) using the delta-Ct method. All RT–qPCR assays were done in triplicate. RNA-sequencing libraries were prepared using TruSeq Total RNA library Prep Kit (Illumina) at Novogene Corporation Inc., and 150bp paired-end sequencing was performed on an Illumina HiSeq 4000 with a depth of 50 million reads per sample. A reference index was generated using GTF annotation from GENCODEv28. Raw FASTQ files were aligned and assembled to GRCh38 and GRCh37 with HISAT2 / Stringtie (V 2.1.0)(5). The differential expression was analyzed in R with packages EdgeR (V 3.32.1) and Rsubread (V1.6.3)(6, 7). Derived counts per million and p-values were used to create a rank ordered list, which was then used for subsequent integrative analysis. Expression levels at known cell type markers from single cell expression databases were used to validate the identity of isolated cell-type populations for methylome analysis(8).

#### Primers used for RT–qPCR in radiation-treated mouse tissues

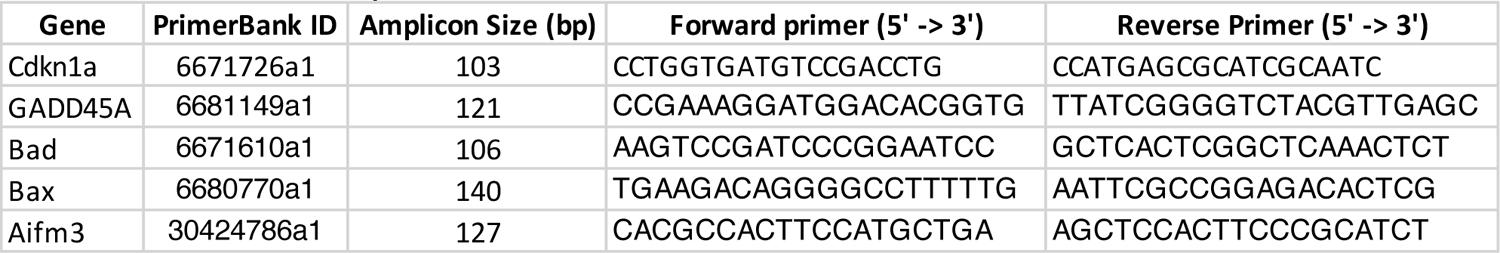

#### Reference DNA methylation data from healthy tissues and cells

Controlled access to reference WGBS data from normal human tissues and cell types was requested from public consortia participating in the International Human Epigenome Consortium (IHEC)(9) and upon approval downloaded from the European Genome-Phenome Archive (EGA), Japanese Genotype-phenotype Archive (JGA), database of Genotypes and Phenotypes (dbGAP), and ENCODE portal data repositories (**Supplemental Table 1**)(10–12). Reference mouse WGBS data from normal tissues and cells were downloaded from selected GEO and SRA datasets (**Supplemental Table 2**)(13–26). Downloaded FASTQs were processed and realigned in a similar manner as the locally generated bisulfite-sequencing libraries described above. However, parameters were adjusted to account for each respective WGBS library type at both trimming and alignment steps as previously described in the Bismark User Guide (http://felixkrueger.github.io/Bismark/Docs/). WBGS libraries were deduplicated using deduplicate_bismark (V 0.22.3) Special consideration of bisulfite conversion efficiency was given to samples prepared by the µWGBS protocol and reads with a bisulfite conversion rate below 90% or with fewer than three cytosines outside a CpG context were removed(27).

#### Identification of cell-type specific methylation blocks

We reduced the original 297 human WGBS samples to a final set of 104 samples to identify differentially methylated cell-type specific blocks. We excluded samples from bulk tissues and those that did not have sufficient coverage (missing values in >50% of methylation blocks). Outlier replicates, or those clustering with fibroblasts or stromal cell types were excluded, due to possible contamination. Only immune cell methylomes that were reprocessed from raw sequencing data to PAT files were used to identify DMBs. We organized the final 104 human reference samples into groupings of 20 cell types (**Supplemental Table 1**). Similarly, the starting 109 mouse WGBS samples were reduced to a final set of 43 samples that were organized into a final grouping of 9 cell types and tissues (**Supplemental Table 2**). Tissue and cell-type specific methylation blocks were identified from reference WGBS data using custom scripts (**Supplemental code**). We performed a one-vs-all comparison to identify differentially methylated blocks unique for each group. This was done separately for human and mouse. From this we first identified blocks covering a minimum of three CpG sites, with length less than 2Kb and at least 10 observations. Then, we calculated the average methylation per block/sample, as the ratio of methylated CpG observations across all sequenced reads from that block. Differential blocks were sorted by the margin of separation, termed “delta beta”, defined as the minimal difference between the average methylation in any sample from the target group vs all other samples. We selected blocks with a delta-beta ≥ 0.4 for human and ≥ 0.35 for mouse. This resulted in a variable number of cell-type specific blocks available for each tissue and cell type. Additional separation of endothelial cell populations from different tissues was performed to identify unique markers for liver endothelial versus cardiopulmonary endothelial blocks that do not overlap. Separately, pan-endothelial blocks were identified with methylation status in common to all endothelial cell populations. For some cell-types, a reduced subset of blocks (ie. top 200) were used for deconvolution in the circulation if the original number identified was greater than one standard deviation above the mean. Selected human and mouse blocks for cell types of interest can be found in **Supplemental Tables 3 and 4.**

#### Methylation score and visualization of cell-type specific methylation atlas

Each DNA fragment was characterized as U (mostly unmethylated), M (mostly methylated) or X (mixed) based on the fraction of methylated CpG sites as previously described(28). We used thresholds of ≤33% methylated CpGs for U reads and ≥66% methylated CpGs for M. We then calculated a methylation score for each identified cell-type specific block based on the proportion of U/X/M reads among all reads. The U proportion was used to define hypomethylated blocks and the M proportion was used to define hypermethylated blocks. Heatmaps were generated using the pretty heatmap function in the RStudio Package for the R bioconductor (RStudioTeam, 2015).

#### In-silico simulations and WGBS deconvolution

In silico mix-in simulations were performed using *wgbstools* (V 0.1.0)(29) to validate the fragment-level deconvolution algorithm at the identified cell-type specific blocks (**Supplemental Figures 3, 5, 6 and 7**). Reference data with greater than three replicates per cell type were split into independent training and testing sets, leaving at least one replicate out for testing. Since the mouse lung endothelial reference WGBS data had only three replicates, sequenced fragments were merged across replicates for this cell type and then randomly split into training (80%) and testing (20%) sets (using wgbstools merge and then wgbstools pat_splitter). The cell-type specific blocks included in the human and mouse methylation atlases were constructed using training set fragments only. For each cell type profiled, we mixed known proportions of target fragments into a background of leukocyte fragments using wgbstools mix_pat. The leukocyte fragments were obtained from n=4 buffy coat samples in mouse and n=5 buffy coat samples in human. We performed three replicates for each admixture ratio assessed (0.05%, 0.1%, 0.5%, 1%, 2%, 5%, 10%, 15%), which were analyzed as described above, and present the average predicted proportion and standard deviation across all replicates. Model accuracy was assessed through correct classification of the actual percent target mixed.

Each mixture was analyzed using our WGBS atlas and fragment-level deconvolution model in contrast to the 450K array atlas and NNLS model described in Moss et al(30). Array based 450K data were simulated using wgbstools beta_to_450k function (V 0.1.0) and deconvolution performed as in Moss et al. (https://github.com/nloyfer/meth_atlas). Our sequencing-based approach allowed for fragment-level cfDNA analysis of CpG methylation patterns, as opposed to relying on the use of single CpG sites from methylation array data(31). From the in-silico mix-in simulations, we found that our probabilistic fragment-level deconvolution model outperforms traditional array-based analysis for each tissue and cell type of interest to validate the prediction accuracy and sensitivity (**Supplemental** Figure 7). We found that pattern analysis at the cell-type specific methylation blocks identified here allowed for accurate detection of cfDNA from a given source when present in less than 0.1% of a mixture, a marked improvement in comparison to current 450K approaches (**Supplemental** Figure 7).

#### Functional annotation and pathway analysis

Cell-type specific methylation blocks were provided as input for analysis in HOMER (V4.11.1) (http://homer.ucsd.edu/homer/)(32). Each block was associated with its closest nearby gene and provided a genomic annotation using the annotatePeaks.pl function, with “-size given -CpG” parameters. By default, TSS (transcription start site) was defined from −1 kb to +100 bp, TTS (transcription termination site) was defined from −100 bp to +1 kb, and CpG islands were defined as a genomic segment with GC content ≥50%, genomic length >200 bp and the ratio of observed/expected CpG number >0.6. Prediction of known and de-novo transcription factor binding motifs were also assessed by HOMER using the findMotifsGenome.pl function. The top 5 motifs based on p-value were selected from each analysis. Pathway analysis of identified tissue and cell-type specific methylation blocks was performed using Ingenuity Pathway Analysis (IPA)(33) (Qiagen) and Genomic Regions Enrichment of Annotations Tool (GREAT)(34). GeneSetCluster was used to cluster identified gene-set pathways based on shared genes(35). Canonical pathways/functional annotations were grouped into clusters by calculating the similarity of pathways/annotations using the relative risk (RR) of each pathway appearing based on the genes enriched within the pathway. RR scores were clustered into groups using kmeans. Over-representation analysis was implemented in the WebgestaltR (ORAperGeneSet) plugin to interpret and functionally label identified gene-set clusters(36). Integration of methylome and transcriptome data generated from tissue-specific endothelial cells was performed using an expanded set of cell-type specific blocks (--bg.quant 0.2) compared to the more restricted set of blocks used for deconvolution analysis in the circulation (--bg.quant 0.1). The extended endothelial-specific methylation blocks can be found in **Supplemental Table 9**.

---

**Supplemental Figure 1.**
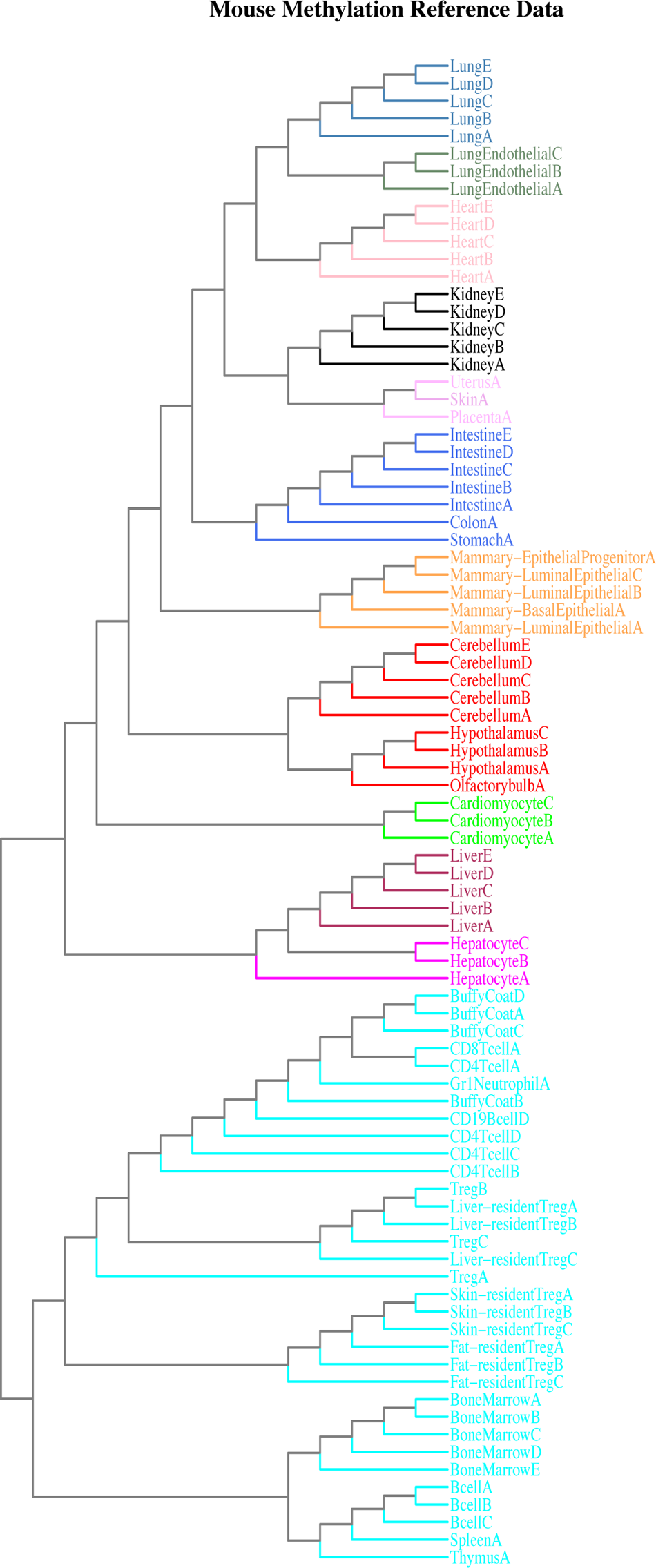
Characterization of mouse cell-type specific reference methylation data. Tree dendrogram depicting relationship between mouse reference Whole Genome Bisulfite Sequencing (WGBS) datasets from different tissues and cell types included in the analysis. Methylation status of the top 30,000 variable blocks was used as input data for the unsupervised hierarchical clustering.

**Supplemental Figure 2.**
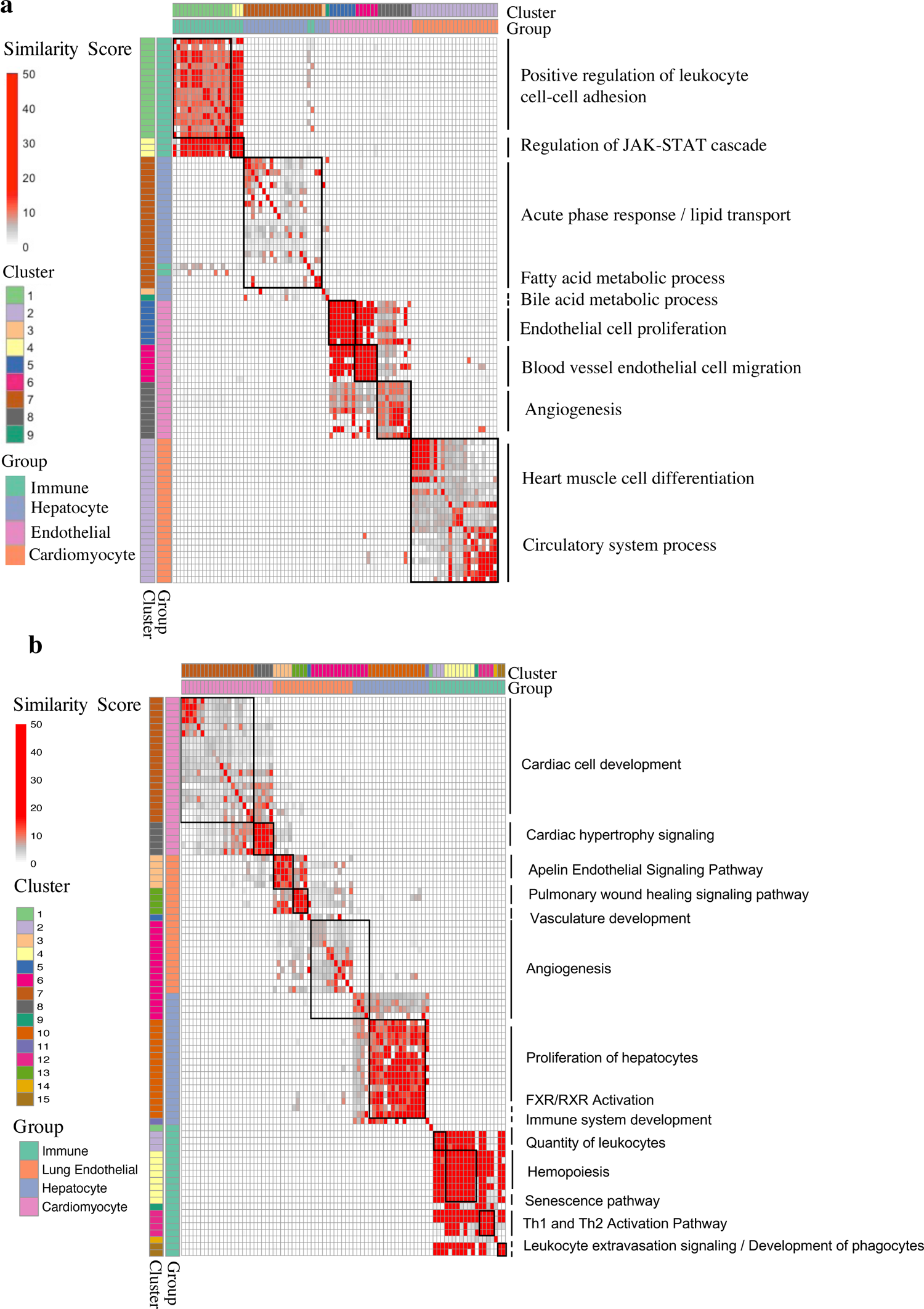
Biological validation of cell-type specific DNA methylation blocks in human and mouse. (a, b) Heatmap of distance scores between gene-set pathways identified from GeneSetCluster. Genes adjacent to human (a) and mouse (b) cell-type specific methylation blocks were identified using HOMER and pathway analysis was performed using both Ingenuity Pathway Analysis (IPA) and GREAT. Significantly enriched gene-set pathways (p<0.05) from differentially methylated blocks identified in immune, cardiomyocyte, hepatocyte, and endothelial cell types were analyzed using GeneSetCluster. Cluster analysis was performed to determine the distance between all identified gene-set pathways based on the degree of overlapping genes from each individual gene set compared to all others. Over-representation analysis was implemented in the WebgestaltR (ORAperGeneSet) plugin to interpret and functionally label identified gene-set clusters.

**Supplemental Figure 3.**
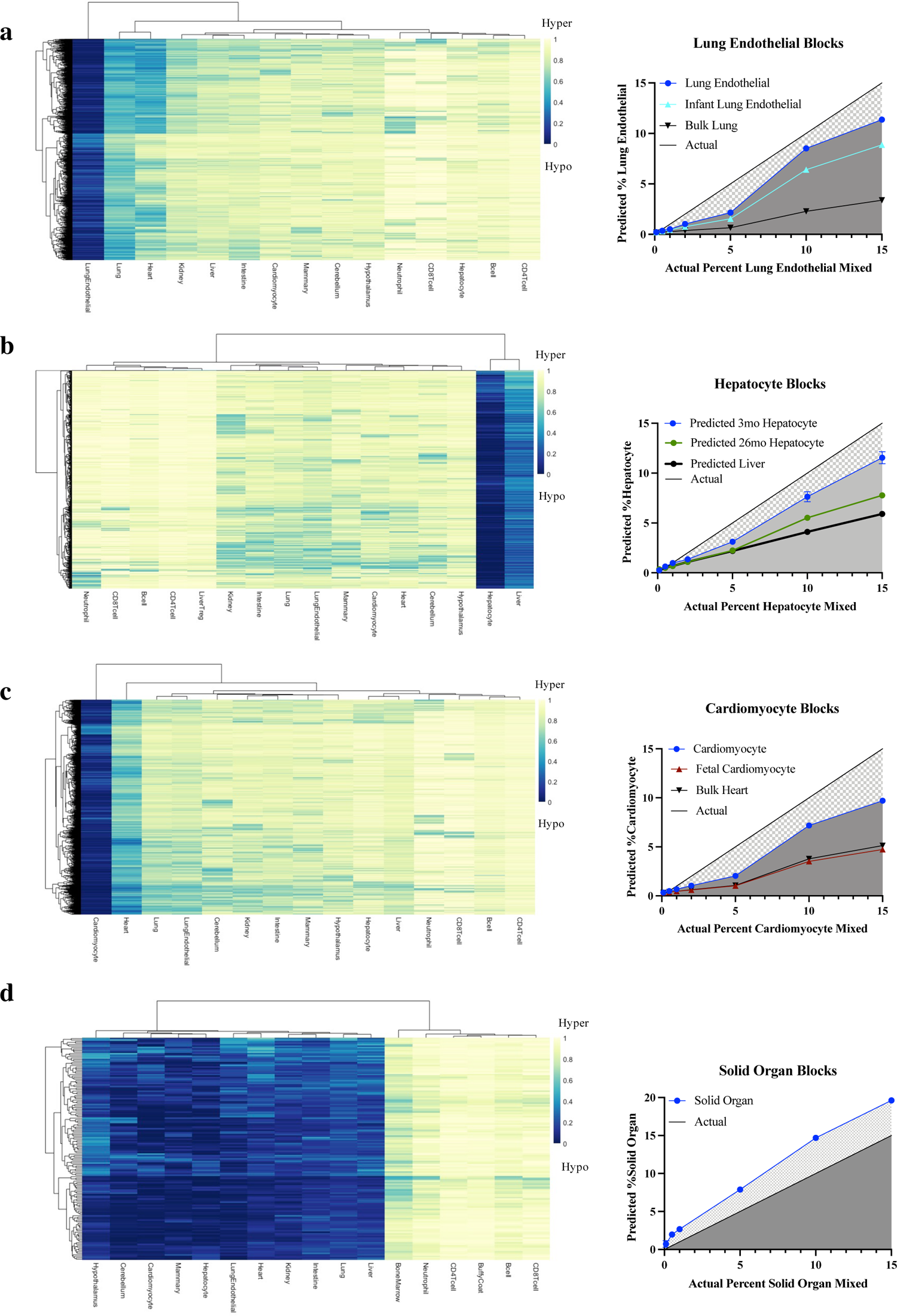
Sensitivity and specificity of identified mouse cell-type specific differentially methylated blocks. (a-d) (Left) Heatmap of all cell-type specific methylation blocks selected for each target cell type. All blocks contain ≥3 CpG sites and have a margin of beta difference greater than or equal to 0.35 separating the target cell type from all others included in the reference maps. All identified methylation blocks for lung endothelial (n=1,546), hepatocyte (n=616), and cardiomyocyte (n=2,917) mouse cell types were hypomethylated. In contrast, all identified immune cell-specific blocks (n=148) were hypermethylated relative to other solid organ cell types in mouse. (Right) In-silico mix-in validation using a fragment-level probabilistic deconvolution model. Target cell-type read-pairs were in-silico mixed into a background of lymphocyte or buffy coat read-pairs at various known percentages (0.5%, 1%, 2%, 5%, 10%, 15%). The deconvolution model was validated on these in-silico mixed samples of known cell-type proportions at the blocks selected. The average predicted %target is shown relative to the known %mixed to assess sensitivity and specificity of the identified cell-type specific blocks and deconvolution model. Data presented as mean ± SD; N=3 replicates per proportion. Reference WGBS samples with less than 3 replicates were split into “0.8 train” to select methylation blocks and “0.2 test” to generate in-silico mixed samples. When available, in-silico mixed samples of the same cell type derived from differently aged mice were also tested (infant < 6 weeks old). In addition, bulk tissue containing the respective cell type was tested as well.

**Supplemental Figure 4.**
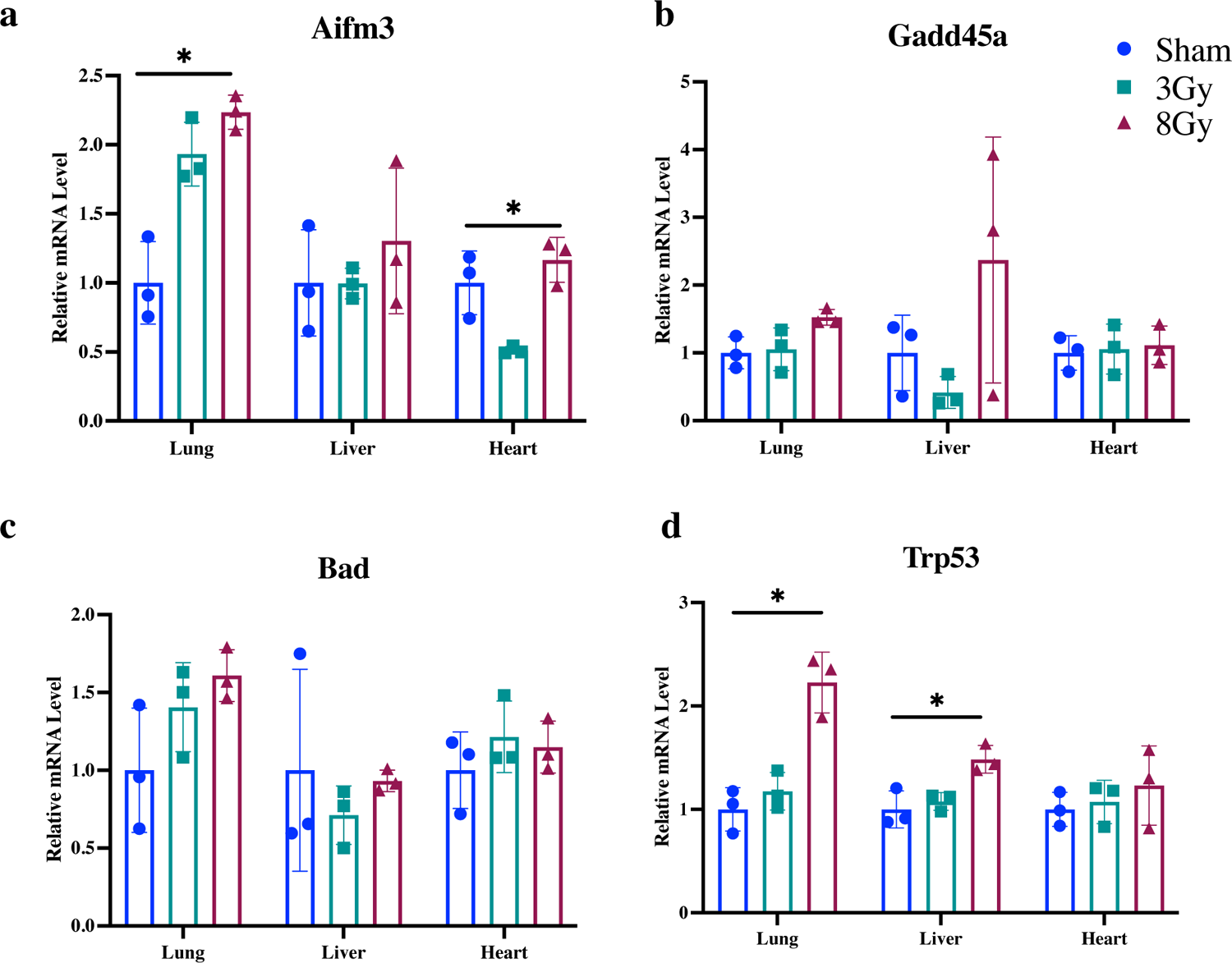
Indicators of damage from radiation in mouse tissues. qPCR analysis of markers of apoptosis and radiation damage (Trp53, Gadd45a, Aifm3, and Bad) in mouse lung, heart, and liver tissues treated with 3Gy and 8Gy radiation compared to sham control. The gene expression in each sample was normalized to the expression of ACTB (beta-actin). Data presented as mean ± SD; N = 3. Kruskal-Wallis test was used for comparisons amongst groups and results were considered significant when *p < 0.05; ns, p ≥ 0.05.

**Supplemental Figure 5.**
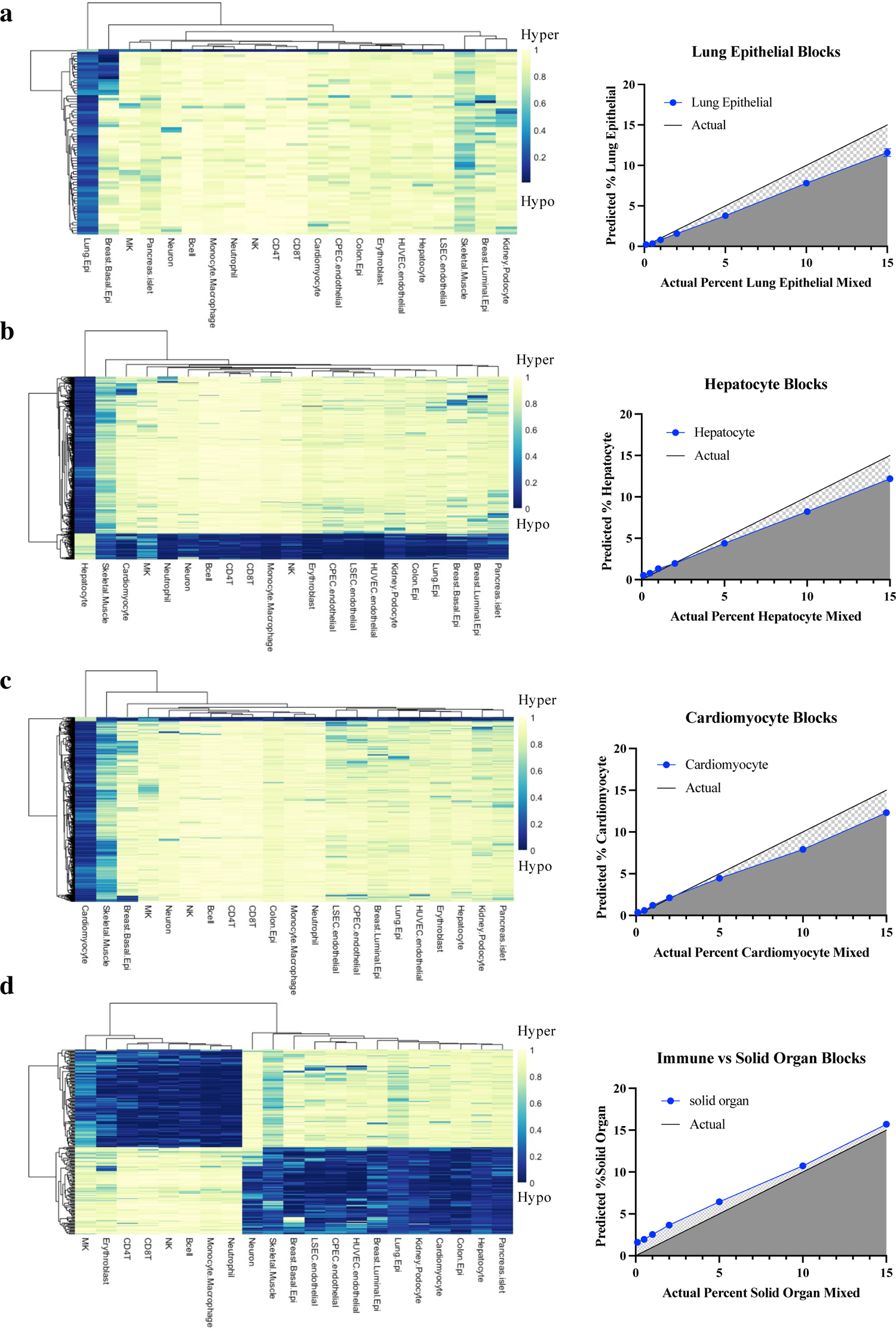
Sensitivity and specificity of identified human cell-type specific differentially methylated blocks. (a-d) (Left) Heatmap of all cell-type specific methylation blocks selected for each target cell type. All blocks contain ≥3 CpG sites and have a margin of beta difference greater than or equal to 0.4 separating the target cell type from all others included in the reference maps. (Right) In-silico mix-in validation from the fragment-level probabilistic deconvolution model. Target cell-type read-pairs were in-silico mixed into a background of lymphocyte or buffy coat read-pairs at various known percentages (0.1%, 0.5%, 1%, 2%, 5%, 10%, 15%). The deconvolution model was validated on these in-silico mixed samples of known cell-type proportions at the blocks selected. The average predicted %target is graphed relative to the known %mixed to assess sensitivity and specificity of the identified cell-type specific blocks and deconvolution model. Data presented as mean ± SD; N=3 replicates per proportion.

**Supplemental Figure 6.**
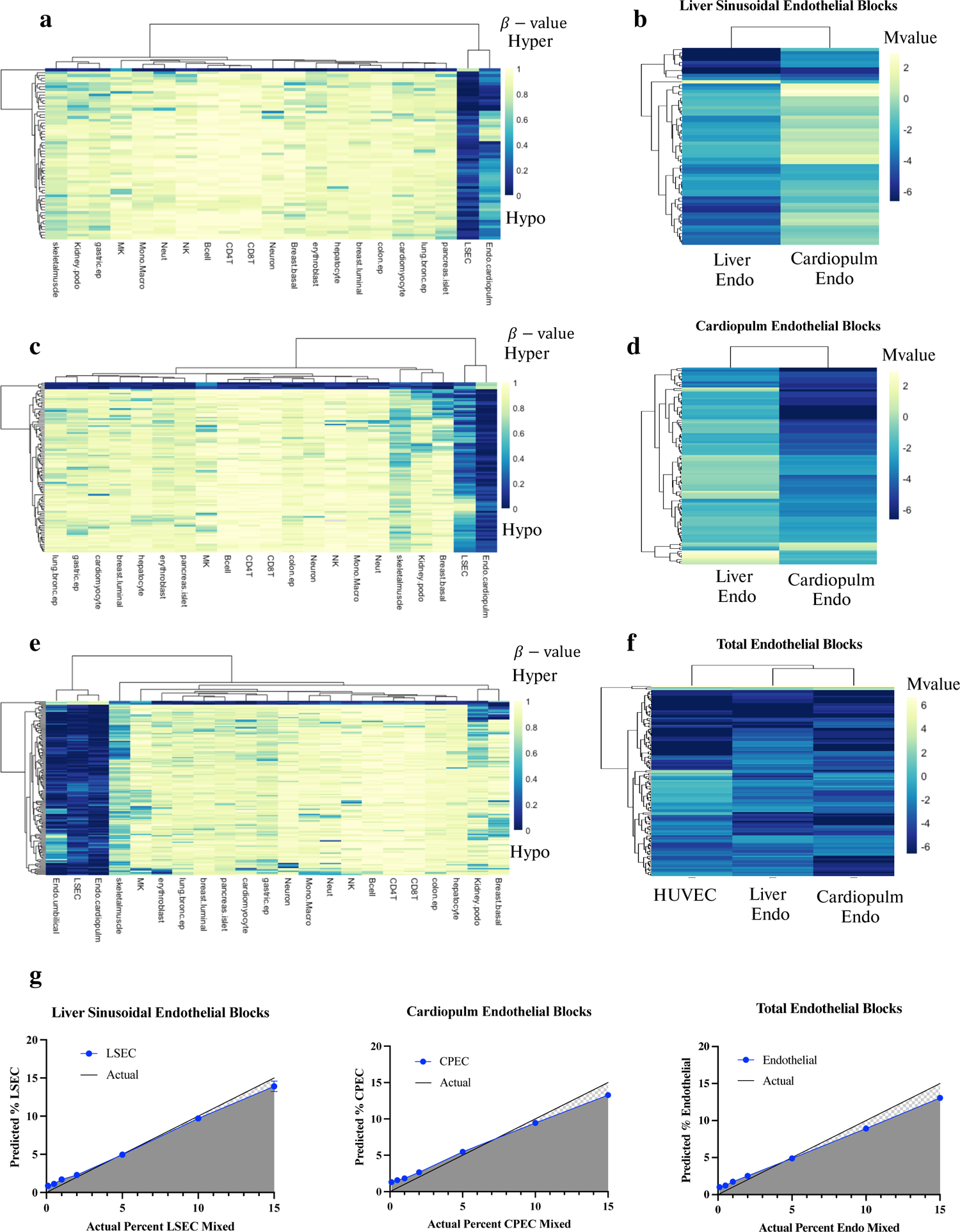
Sensitivity and specificity of identified human endothelial specific differentially methylated blocks. (a, c, e) Heatmap of cell-type specific methylation blocks selected for each target cell type. All blocks contain ≥3 CpG sites and have a margin of beta difference greater than or equal to 0.4 separating the target cell type from all others included in the reference maps. (b, d, f) Enlarged heatmap of cardiopulmonary (b) or liver endothelial (d) specific methylation blocks that are unique relative to other liver or cardiopulmonary endothelial blocks respectively; (f) pan-endothelial specific methylation blocks with common methylation status amongst cardiopulmonary, liver, and HUVEC endothelial cell populations. Methylation status is represented by M-values (Logit transformation of β-values) to limit heteroscedasticity in visual representation of methylation differences across regions. (g) In-silico mix-in validation from a fragment-level probabilistic deconvolution model. Target cell-type read-pairs were in-silico mixed into a background of lymphocyte or buffy coat read-pairs at various known percentages (0.1%, 0.5%, 1%, 2%, 5%, 10%, 15%). The deconvolution model was validated on these in-silico mixed samples of known cell-type proportions at the blocks selected. The average predicted %target is graphed relative to the known %mixed to assess sensitivity and specificity of the identified cell-type specific blocks and deconvolution model. Data presented as mean ± SD; N=3 replicates per proportion.

**Supplemental Figure 7.**
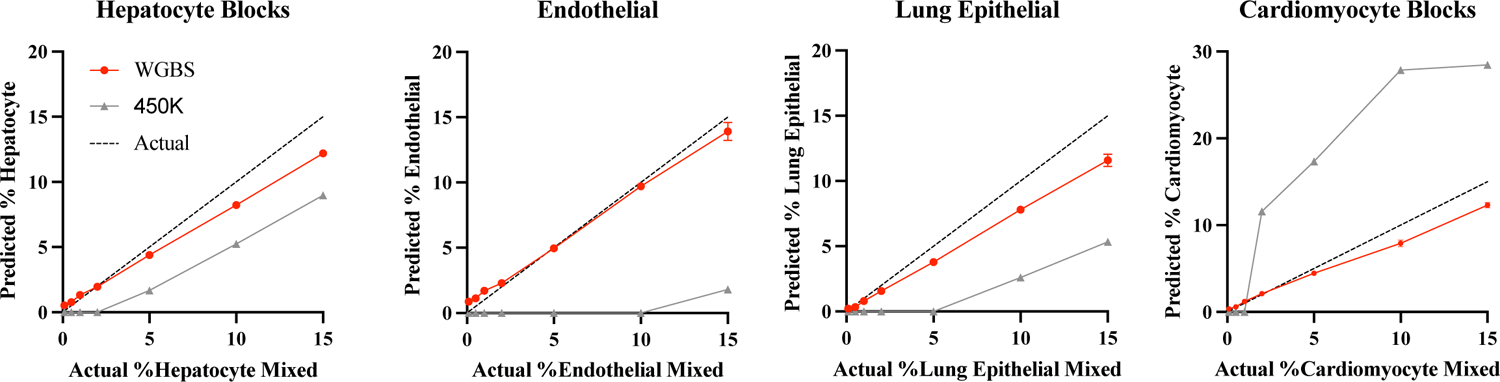
Performance of the probabilistic fragment-level deconvolution algorithm using WGBS data relative to NNLS MethAtlas from 450K array data. Cell type-specific markers outperform the array-based atlas and achieve <0.1% resolution. Shown are in silico simulations for four cell types, computationally mixed within leukocytes at various known percentages (0.05%, 0.1%, 0.5%, 1%, 2%, 5%, 10%, 15%). Each mixture was analyzed using our WGBS atlas and fragment-level deconvolution model (red), compared to Moss et al. 2018 (gray). Data presented as mean ± SD; N=10 replicates per proportion.

**Supplemental Figure 8.**
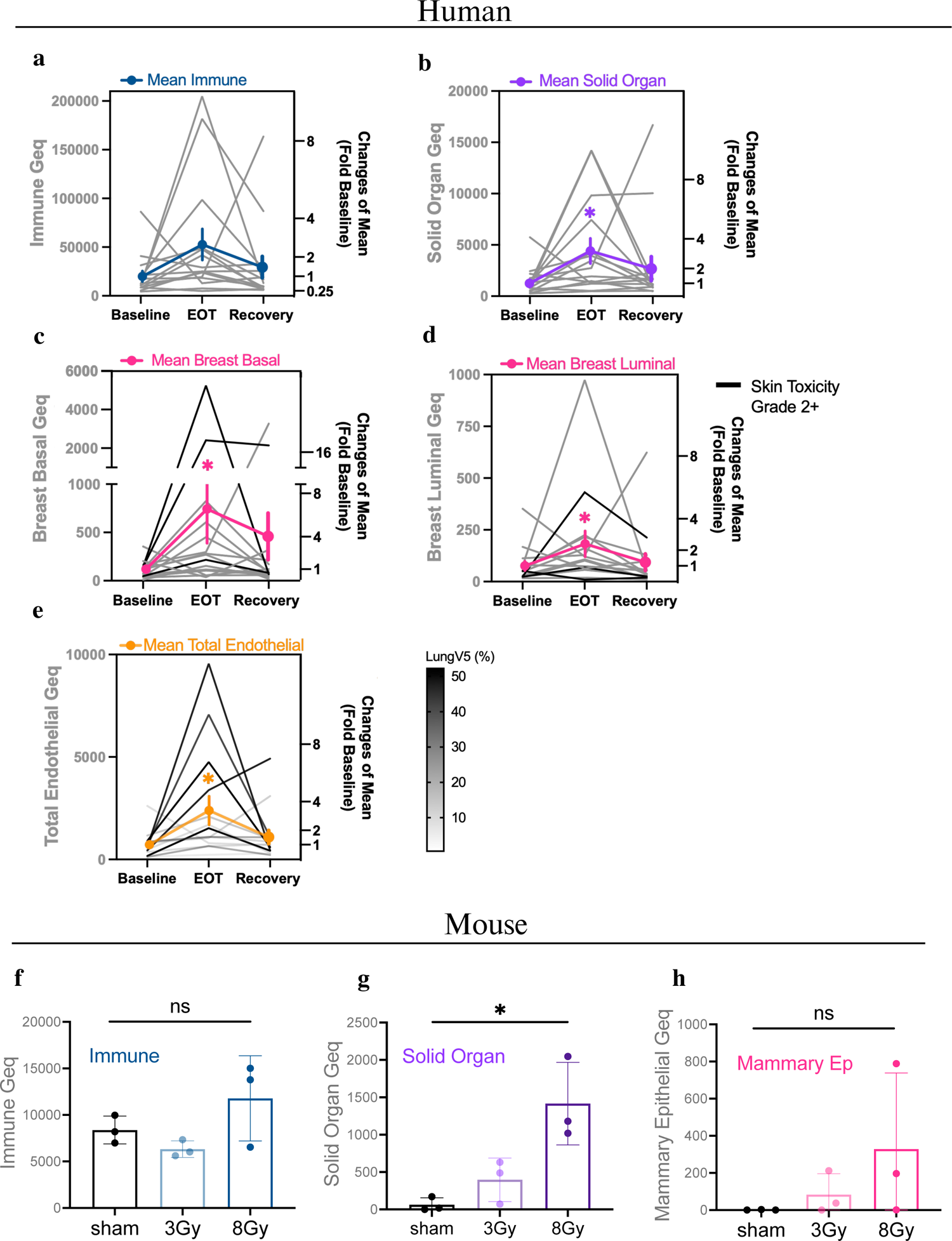
Radiation-induced effects on immune and other solid organ cell-types in human (a-e) and mouse (f-h) samples. (a) Predicted human immune-derived cfDNA in Geq. Human Geq are calculated by multiplying the relative fraction of cell-type specific cfDNA x initial concentration cfDNA ng/mL x the weight of the haploid human genome. Immune cfDNA was assessed at n = 222 methylation blocks found to separate immune cell types from solid organ cell types. (immune = Bcell, CD4Tcell, CD8Tcell, NK, MK, Erythroblast, Monocyte, Macrophage, Neutrophil; solid organ = breast basal/luminal epi, lung epi, hepatocyte, kidney podocyte, pancreas islet, colon epi, cardiomyocyte, LSEC, CPEC, HUVEC, neuron, and skeletal muscle). (b) Predicted human solid organ-derived cfDNA in Geq where %solid organ is defined as 100-%immune using the above n=222 methylation blocks. (c) Breast basal epithelial cfDNA (in Geq/mL). Fragment-level deconvolution used top breast basal epithelial specific methylation blocks (n=200). (d) Breast luminal epithelial cfDNA (in Geq/mL). Fragment-level deconvolution used breast luminal epithelial specific methylation blocks (n=330). (e) Predicted total endothelial cfDNA (in Geq/mL). Fragment-level deconvolution was assessed at n = 131 methylation blocks found to separate endothelial cells from all other cell types. (endothelial = CPEC, LSEC, HUVEC; non-endothelial = Bcell, CD4Tcell, CD8Tcell, NK, MK, Erythroblast, Monocyte, Macrophage, Neutrophil, breast basal/luminal epi, lung epi, hepatocyte, kidney podocyte, pancreas islet, colon epi, cardiomyocyte, neuron, and skeletal muscle). (a-e) Friedman test was performed for comparisons amongst groups. ns, p ≥ 0.05; *, p < 0.05; immune p=0.07, solid organ p=0.008, breast basal epithelial p=0.002, breast luminal epithelial p=0.02, total endothelial p=0.01. Mean fold change relative to baseline is presented as mean ± SEM; N = 15. (f) Predicted mouse immune-derived cfDNA in Geq. Mouse Geq are calculated by multiplying the relative fraction of cell-type specific cfDNA x initial concentration cfDNA ng/mL x the weight of the haploid mouse genome. Immune cfDNA was assessed at n = 148 methylation blocks found to separate immune cell types from solid organ cell types. (immune = Bcell, CD4Tcell, CD8Tcell, Neutrophil; solid organ = mammary epi, cardiomyocyte, hepatocyte, lung endothelial, cerebellum, hypothalamus, colon, intestine, kidney). (g) Predicted mouse solid organ-derived cfDNA (in Geq/mL). (h) Mammary epithelial cfDNA (in Geq/mL). Fragment-level deconvolution used mouse mammary epithelial specific methylation blocks (n=874). (f-h) Mean ± SD; N = 3 independent methylome preparations. Kruskal-Wallis test was used for comparisons amongst groups. ns, p ≥ 0.05; *, p < 0.05; immune p = 0.20, solid organ p = 0.01, mammary epithelial p=0.19.

**Supplemental Figure 9.**
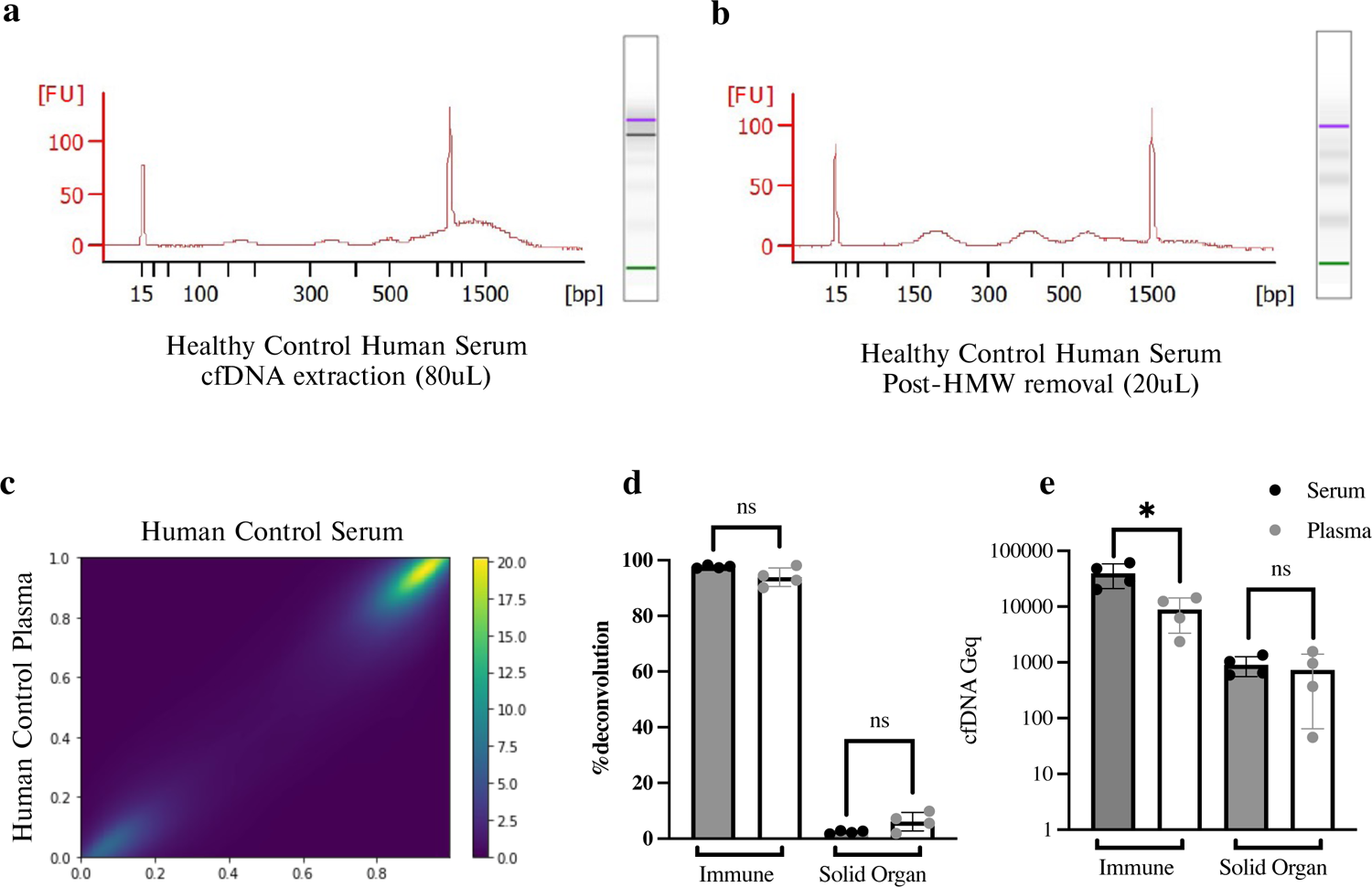
Comparison of methylation status and cellular origins of cfDNA isolated from serum and plasma of healthy human controls. (a, b) Representative bioanalyzer trace of freshly isolated cfDNA extracted from healthy control human serum before (a) and after (b) removal of high-molecular weight (HMW) DNA. (c) Density scatter plot comparing methylation status across blocks in cfDNA isolated from control human serum (n=4) versus control human plasma (n=4). Methylation levels are highly correlated at the block-level with Pearson’s r =0.95 and R^2^=0.89. (d) Predicted %Immune versus %Solid Organ derived cfDNA extracted from either serum or plasma. Origins were assessed at n = 222 methylation blocks found to separate immune cell types from solid organ cell types. (e) Immune and solid organ Geq from cfDNA isolated from serum versus plasma. (d, e) Data presented as mean ± SD; N=4 samples per group. Mann-Whitney test was used for comparisons amongst groups. ns, p ≥0.05; *, p < 0.05.

## Legends for Supplementary Data File 1

**Supplemental Table S1.** Human reference methylation data from healthy tissues and cell types.

**Supplemental Table S2.** Mouse reference methylation data from healthy tissues and cell types.

**Supplemental Table S3.** Identified human cell-type specific methylation blocks. Annotation was performed using Homer. The margin of separation represents the delta-beta (maximum higher – minimum lower) across all samples. Blocks with a (-) direction are hypomethylated and (+) direction are hypermethylated. AMF (average methylation fraction) indicated as a fraction.

**Supplemental Table S4.** Identified mouse tissue and cell-type specific methylation blocks. Annotation was performed using Homer. The margin of separation represents the delta-beta (maximum higher – minimum lower) across all samples. Blocks with a (-) direction are hypomethylated and (+) direction are hypermethylated. AMF (average methylation fraction) indicated as a fraction.

**Supplemental Table S5.** Summary of identified human (A) and mouse (B) cell-type specific methylation blocks.

**Supplemental Table S6.** Mouse cfDNA sample concentrations and predicted cell-type proportions from deconvolution analysis at identified cell-type specific blocks for target cell types.

**Supplemental Table S7.** Human cfDNA sample concentrations and predicted cell-type proportions from deconvolution analysis at identified cell-type specific blocks for target cell types.

**Supplemental Table S8.** Characteristics of breast cancer patients enrolled in this study.

**Supplemental Table S9.** Extended endothelial-specific methylation blocks (bg.quant 0.2) used for pathway analysis and validation of cell identity through integration with paired RNA expression data.

**Supplemental Table S10.** Genomic annotation of identified human and mouse cell-type specific hypomethylated and hypermethylated blocks relative to all captured blocks.

**Supplemental Table S11.** Top 25 significantly enriched biological pathways and functions for genes associated with differential methylation in each cell-type.

**Supplemental Table S12.** Extended clinical data and characteristics of breast cancer patients enrolled in this study.

